# Reducing growth-medium complexity reveals nutrient-responsive programs in a near-minimal bacterium

**DOI:** 10.64898/2026.09.29.755377

**Authors:** Anthony Duval, Jérémy Gagnon, Simon Jeanneau, Dominick Matteau, Pierre-Étienne Jacques, Sébastien Rodrigue

## Abstract

*Mesoplasma florum* is a fast-growing, near-minimal bacterium and an emerging model for systems and synthetic biology. However, its dependence on complex serum-containing media limits experimental control and complicates the interpretation of cellular phenotypes. Here, we developed CMRL-AT, a serum-free, quasi-defined medium that supports rapid growth comparable to the commonly used ATCC 1161 medium. Despite supporting similar biomass, CMRL-AT profoundly reshaped the *M. florum* transcriptome, with approximately one-third of the annotated protein-encoding genes being differentially expressed relative to ATCC 1161. These changes revealed distinct physiological programs associated with rapid growth in complex medium and higher nutrient acquisition in CMRL-AT, illustrating how medium composition alters the functional priorities of a near-minimal cell. Transcriptome profiling across six energy sources further uncovered distinct sugar-responsive expression programs. Combining these responses with transcription-unit organization, protein-domain predictions, and metabolic context resolved fructose- and sucrose-responsive modules, and allowed the assignment of previously ambiguous phosphotransferase system components to specific sugar-utilization pathways. CMRL-AT provides an experimental framework to help resolving gene functions, refining metabolic models, and designing reduced genomes adapted to defined environments.

## Introduction

Advances in DNA synthesis, assembly, and sequencing technologies have greatly expanded our ability to engineer biological systems. DNA fragments spanning thousands of base pairs can now be synthesized and sequence-verified rapidly and at relatively low cost, facilitating the construction of synthetic chromosomes and entire genomes, an approach broadly referred to as synthetic genomics (Fu & Shen, 2024). Key achievements in this field include the creation of JCVI-syn1.0 and JCVI-syn3.0, synthetic bacteria derived from *Mycoplasma mycoides* subsp. *capri*, the latter containing a highly reduced genome encoding only 473 genes (Hutchison *et al*, 2016). These advances have highlighted both the potential of genome-scale engineering and the substantial gaps that remain in our understanding of the fundamental principles governing cellular life.

Despite these achievements, synthetic genomics remains largely constrained to top-down approaches that modify existing genomes through reduction, recoding, or refactoring. The design of truly de novo genomes remains difficult because the fundamental principles governing genome organization and cellular function are still incompletely understood. The challenge is evident even in *Escherichia coli* MG1655, one of the best-characterized organisms, where approximately one-third of genes still lack a clear functional assignment (Karp *et al*, 2025; Moore *et al*, 2024), limiting our ability to build predictive cellular models (Karr *et al*, 2012). Resolving gene function depends on interpreting cellular behavior under controlled conditions, which in turn requires growth media of defined composition. Consequently, many researchers have turned to the Mollicutes, which include some of the simplest autonomously replicating cells known, as experimental systems for uncovering the minimal requirements for cellular life (Glass *et al*, 2017; Morowitz, 1984).

Mollicutes are a class of bacteria characterized by small cell sizes (∼0.2 to 0.8 μm) and reduced genomes ranging from approximately 580 to 2,200 kb. Extensive genome reduction during their evolution has resulted in the loss of numerous metabolic functions, including cell wall synthesis (Sirand-Pugnet *et al*, 2007). Although Mollicutes are best known for containing numerous medically and agriculturally important pathogens, their genomic simplicity has also made them attractive model organisms for systems and synthetic biology research (Matteau *et al*, 2024). The growing availability of omics datasets, genome-scale models, and genome engineering tools has enabled detailed investigations of minimal cellular life and the development of simplified biological chassis for biotechnology. Notable examples include the construction and characterization of the JCVI-syn3A minimal cell derived from *Mycoplasma mycoides* subsp. *capri* (Gibson *et al*, 2010; Hutchison *et al*, 2016; Breuer *et al*, 2019), the engineering of *M. pneumoniae* for therapeutic applications (Matteau & Rodrigue, 2021; Garrido *et al*, 2021), and the development of whole-genome engineering platforms in Mollicutes species (Talenton *et al*, 2022).

Closely related to mycoplasmas, *Mesoplasma florum* has emerged as an attractive model organism for systems biology and synthetic genomics (Matteau *et al*, 2024). In contrast to many Mollicutes of medical or veterinary importance, *M. florum* has not been associated with disease and is instead thought to be a commensal of arthropods (Matteau *et al*, 2024). With a 793-kb genome encoding approximately 720 genes, the L1 type strain is among the smallest autonomously replicating bacteria currently characterized (GenBank: AE017263.1). *M. florum* also exhibits one of the fastest growth rates among Mollicutes, with a doubling time of approximately 32 min in rich medium (Matteau *et al*, 2020). Recent work has generated an extensive toolkit for *M. florum*, including genetic manipulation methods, whole-genome engineering approaches, comparative genomics resources, transposon mutagenesis datasets, genome-scale metabolic models, and integrative cellular characterizations (Baby *et al*, 2018b, 2018a; Matteau *et al*, 2017). Together, these resources have established *M. florum* as a powerful platform for investigating minimal cellular life. Proposed minimal genome scenarios have further revealed important differences relative to JCVI *M. mycoides* synthetic cells, highlighting the plasticity of minimal gene sets and motivating efforts toward the construction and study of minimized *M. florum* genomes (Baby *et al*, 2018b).

Like other Mollicutes, *M. florum* requires a nutrient-rich medium to compensate for its extensive metabolic deficiencies in vitro (Matteau *et al*, 2024). The medium most commonly used for its cultivation is ATCC 1161, a complex formulation containing horse serum, yeast extract, and heart infusion broth (Matteau *et al*, 2017). Because these biologically derived ingredients vary between batches, they can introduce substantial experimental variability. In contrast, defined media provide precise control over nutrient availability and facilitate systematic investigations of cellular physiology. Such media are valuable for omics analyses, functional genomics studies, and metabolic modeling efforts, as they allow experimental perturbations to be interpreted within a well-defined nutritional context (diCenzo *et al*, 2018; Jensen *et al*, 2020; Sastry *et al*, 2021; Gaspari *et al*, 2020). Previous efforts to simplify the growth requirements of *M. florum* led to the development of the CSY medium and enabled validation of predictions from its genome-scale metabolic model (Lachance *et al*, 2021). CSY uses CMRL-1066, a chemically defined mammalian cell culture medium that has also served as the basis of formulations for other Mollicutes species (Burgos *et al*, 2023; Gardella & Del Giudice, 1995). However, complete elimination of serum and yeast extract was not possible, as reducing their concentrations substantially *impaired M. florum* growth (Lachance *et al*, 2021). Thus, a robust serum-free medium capable of supporting rapid and reproducible growth of *M. florum* remains to be developed.

Here, we describe the development of CMRL-AT, a serum-free growth medium that supports rapid growth of *M. florum* under substantially more defined conditions than conventional media. We reasoned that removing serum and yeast extract would expose the metabolic dependencies these ingredients normally mask, and that the resulting transcriptional response would identify the pathways *M. florum* recruits to compensate. Using RNA-seq and a refined genome annotation, we characterized this response and interpreted it in the context of the previously published genome-scale metabolic model iJL208 (Lachance *et al*, 2021). Defined conditions further allowed us to vary carbon sources and resolve the sugar transport and catabolic pathways that remained ambiguous in the current annotation. Together, these results provide new insights into the physiological adaptation of *M. florum* to defined growth conditions and further strengthen its utility as a model system for systems and synthetic biology.

## Results

### Development of a serum-free growth medium for *M. florum*

To develop a serum-free medium for *M. florum*, we supplemented the CMRL-1066 mammalian cell culture medium with varying concentrations of tryptone and Albumax II. Supplementation with either component alone promoted growth in a dose-dependent manner (Fig. S1A,B). Combined supplementation showed a similar trend, with cell densities plateauing at 15 mg/mL of each supplement (Fig. S1C). Higher concentrations increased background growth in the absence of sugar (Fig. S1D). Accordingly, 15 mg/mL Albumax II and 15 mg/mL tryptone were selected for subsequent experiments since they maximized growth while maintaining minimal sugar-independent background growth (Fig. S1C,D).

To determine the optimal sugar concentration for the resulting medium, we performed dose-response experiments using six carbohydrates previously shown to support *M. florum* growth in CSY medium (Lachance *et al*, 2021). For glucose, maximal cell densities after 24 h were obtained at 6% supplementation, without evidence of growth inhibition after 48 h (Fig. S2A). Similar results were observed for all tested sugars, including a mixture containing equal concentrations of all six carbohydrates (Fig. S2A-G). Therefore, all subsequent CMRL-AT formulations contained sugars at a final concentration of 6%.

Under these optimized conditions, CMRL-AT supported growth approaching that observed in the complex ATCC 1161 medium (Fig. 1B,C and Fig. S3,S4). Depending on the supplied carbohydrate, *M. florum* exhibited doubling times ranging from approximately 42 to 54 min, with glucose supporting the fastest growth among individual sugars tested (Fig. 1B and Fig. S3). A formulation containing 1% of each sugar yielded the shortest doubling time (∼40 min). In comparison, cultures grown in ATCC 1161 displayed a doubling time of approximately 31 min, whereas CSY supplemented with 6% sucrose exhibited a substantially slower doubling time of approximately 70 min (Fig. 1B and Fig. S3). CMRL-AT supplemented with 6% glucose also achieved cell densities comparable to those observed in ATCC 1161, reaching maximal biomasses of 3.4 × 10⁹ and 4.2 × 10⁹ cells mL⁻¹, respectively (Fig. 1C and Fig. S4). By contrast, CSY supplemented with 6% sucrose reached only 3.2 × 10⁸ cells mL⁻¹ under the same growth conditions (Fig. 1C and Fig. S4).

**Figure 1.**
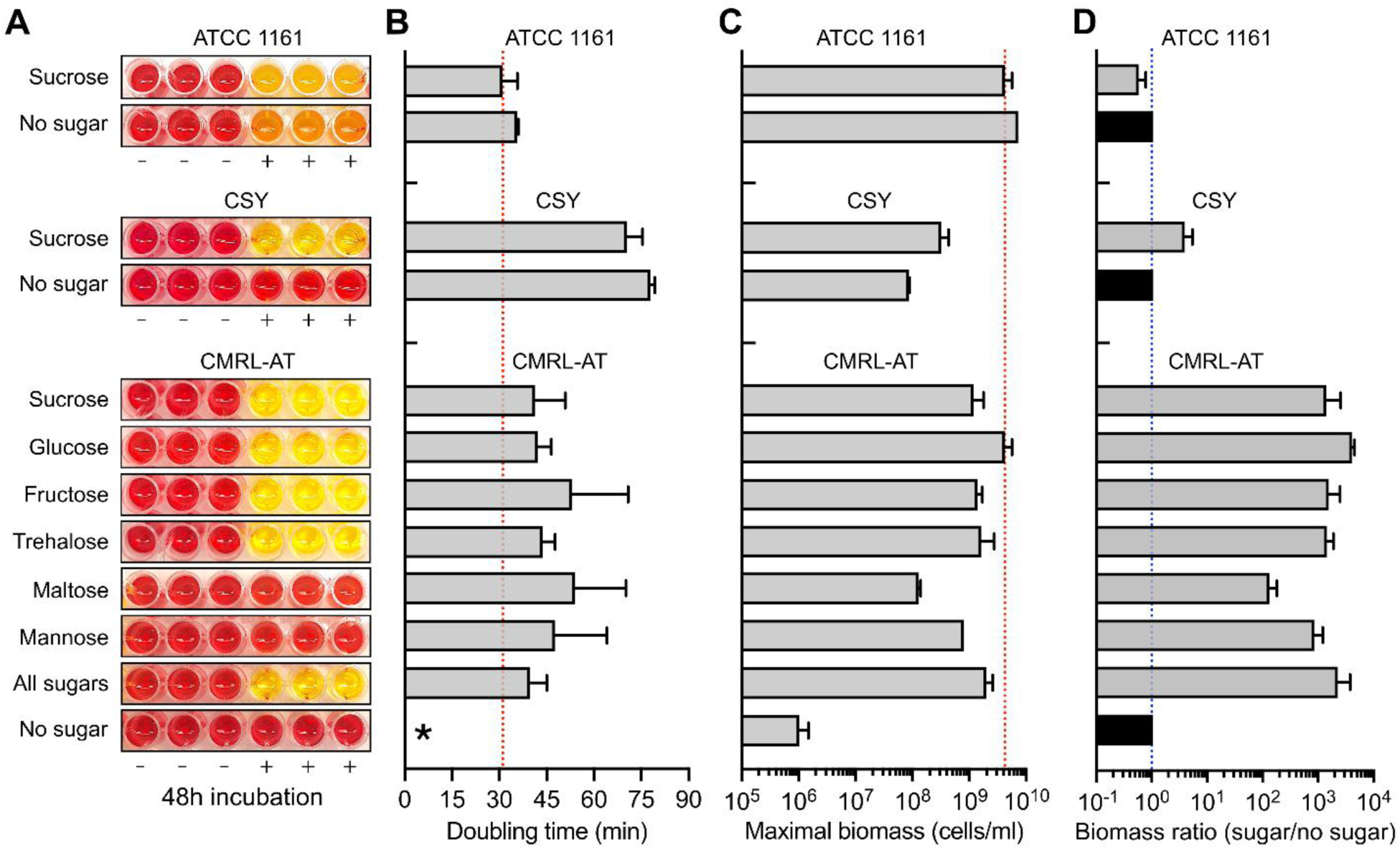
Analysis of *Mesoplasma florum* growth in three different media. **A)** Bacterial growth assay using culture medium color change due to acidification in three different media (ATCC 1161, CSY, and the defined CMRL-AT), supplemented or not with various sugars. Sugars were supplemented at 4% in ATCC 1161, and at 6% in CSY and CMRL-AT. (+), inoculated wells; (–) non-inoculated controls. The picture was taken after a 48h incubation period. **B-C)** Doubling time (B) and maximal biomass (C) of *M. florum* L1 in the three different media, with or without sugar supplementation, determined by colony-forming units (CFUs). The red dotted lines indicate the values obtained for the ATCC 1161 medium with 4% sucrose. Asterisks mark non-calculable doubling time due to insufficient growth. See Figure S3 for complete growth curves. **D)** Biomass ratio between sugar supplemented and non-supplemented conditions determined for each growth medium. Growth conditions without sugars were arbitrarily set to a ratio of 1, as also denoted by the blue dotted line. Bars and error bars indicate mean and standard deviation calculated over biological duplicates, respectively.

A key advantage of CMRL-AT was its very low background growth in the absence of added sugar (Fig. 1). Whereas *M. florum* reached similar cell densities in ATCC 1161 regardless of sugar supplementation, growth in CMRL-AT was highly restricted when no sugar was added (Fig. 1, Fig. S3 and Fig. S4). Consequently, maximal biomass increased by 133-fold (maltose) to 4,134-fold (glucose) upon sugar supplementation in CMRL-AT, while supplementation of CSY with 6% sucrose increased biomass by only approximately fourfold (Fig. 1C). These observations were consistent with changes in medium color after 48 h of incubation (Fig. 1A). In the absence of sugar, CMRL-AT and CSY exhibited little or no color change, whereas ATCC 1161 shifted from orange to yellow regardless of sugar supplementation. Finally, the stationary phase was markedly prolonged in CMRL-AT relative to ATCC 1161 containing 4% sucrose, a phenomenon also observed in CSY and in ATCC 1161 without added sucrose (Fig. S3).

### Transcriptome profiling using a multi-source genome annotation

Although the newly developed CMRL-AT medium provided growth characteristics comparable to those of ATCC 1161 (Fig. 1), important differences remain in their compositions. To investigate how *M. florum* adapts to these environments, we performed RNA-seq on cultures grown in ATCC 1161 and CMRL-AT supplemented with a mixture of six sugars. Because the six-sugar formulation yielded the fastest growth in CMRL-AT (Fig. 1B and Fig. S3), it was also used for transcriptomic comparisons with ATCC 1161. Additional RNA-seq experiments were conducted on CMRL-AT supplemented with each sugar individually, yielding a total of eight growth conditions (Table S1).

To improve the functional interpretation of transcriptomic data, gene expression was quantified using a genome annotation integrating four independent annotation resources: RAST 2018 (Baby *et al*, 2018b); File S1), GenBank 2014 (File S2), RefSeq 2022 (File S3), and PATRIC 2015 (File S4). Comparison of these annotations revealed substantial disagreement in predicted protein-encoding genes (PEGs), with only 599 PEGs shared among all four annotation sources (Fig. 2A). The remaining 186 PEGs were shared by three or fewer annotation sets, including nearly 100 PEGs unique to a single annotation source (Fig. 2A).

**Figure 2.**
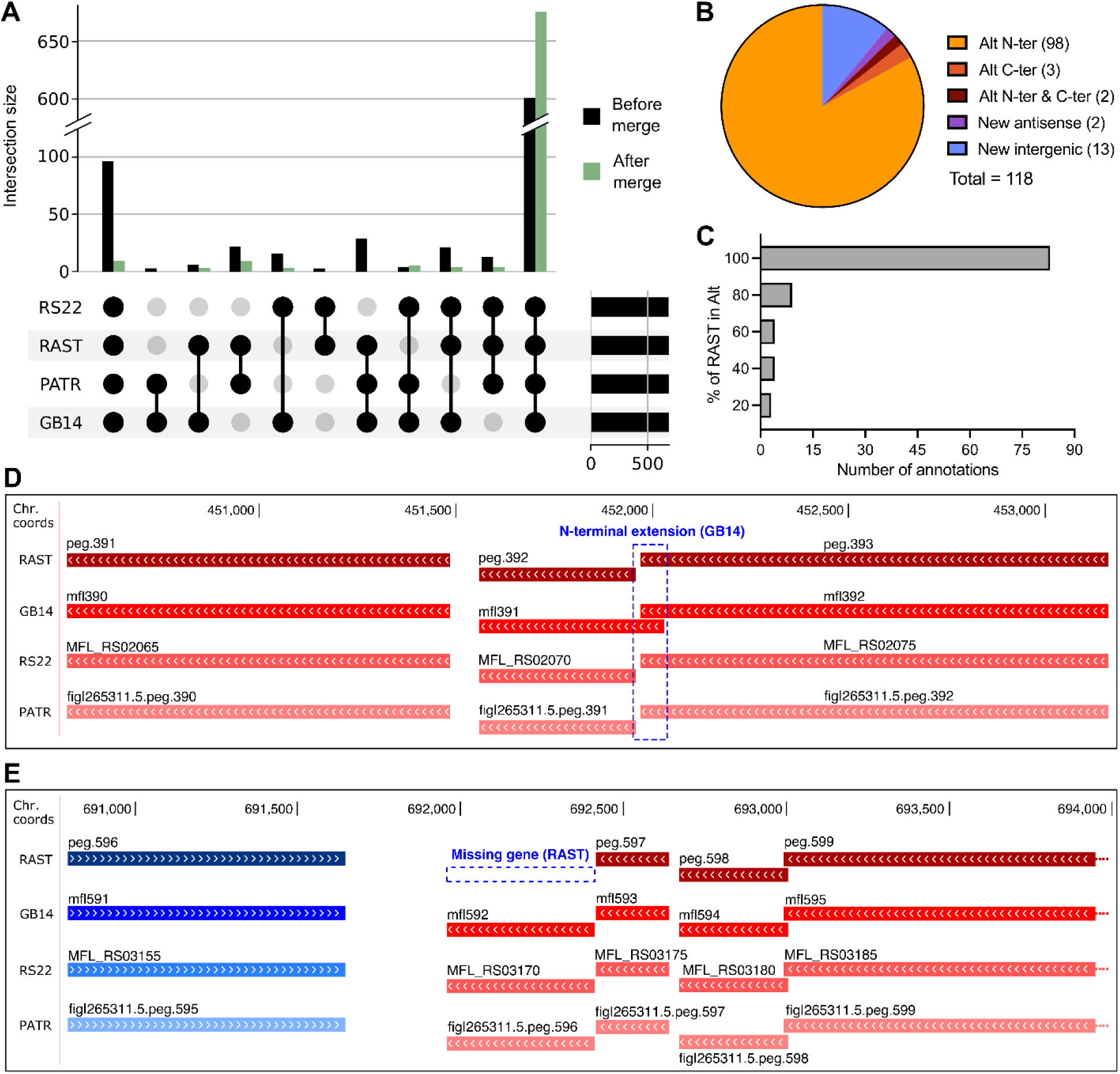
Combination of four different *M. florum* genome annotation sources. **A)** Upset chart showing protein-encoding genes shared with up to four different annotation sources or unique to a single annotation system, before and after the genome annotation merge. Y-axis line break is for representation purposes. RS22, RefSeq update 2022 (NC_006055.1); RAST, RAST 2018 (Baby *et al*, 2018b); PATR, PATRIC update 2015 ([ID:265311.5](https://www.ncbi.nlm.nih.gov/nuccore/NC_006055.1)); GB14, GenBank update 2014 (AE017263). See File S1-S4 for additional information. **B)** Proportion of annotations not identical to RAST annotations, regrouped into five different categories: alternative N-terminal, alternative C-terminal, alternative N and C-terminal, new antisense gene, as well as new intergenic gene. **C)** Distribution of the overlapping proportion of RAST annotations relative to alternative annotations provided by the other annotation sources (see panel B). **D)** Example of an open-reading frame N-terminal extension (*mfl391*) present in at least one annotation source (GB14) relative to the RAST annotation (peg.392). **E)** Example of new intergenic gene added in the enhanced RAST annotation following comparison with the three other annotation sources.

Further examination of these discrepancies showed that most corresponded to alternative translation start sites rather than entirely distinct genes. Specifically, 98 PEGs differed only by their translation start codon relative to the RAST annotation, whereas only three exhibited alternative stop codons and two differed at both the N- and C-termini (Fig. 2B,D and Table S2). Moreover, 81% of these alternative annotations completely encompassed the corresponding RAST feature, indicating putative 5′ or 3′ extensions of existing gene models (Fig. 2C,D and Table S2). Nevertheless, 15 features were identified on the opposite strand or in intergenic regions relative to existing RAST PEGs (Fig. 2B,E).

Following manual curation, overlapping annotations from GenBank, RefSeq, and PATRIC were merged with the RAST annotation to generate a refined multi-source annotation (File S5). This process raised the number of PEGs shared across all annotation sources from 599 to 673 and incorporated nine additional PEGs not represented in the original RAST annotation (Fig. 2A,E and Table S2). Throughout the remainder of this study, genes are referred to using RefSeq 2022 locus tags when available, or otherwise by their GenBank 2014 identifiers.

Using this refined annotation, expression was detected for every annotated *M. florum* PEG across all growth conditions (Fig. S5 and Table S3). Gene expression measurements were highly reproducible, with strong correlations observed among biological replicates within each condition (Fig. S6A). High correlations were also observed across most conditions (R² > 0.91), with the notable exceptions of ATCC 1161 supplemented with six sugars (A6S) and CMRL-AT supplemented with mannose (Fig. S6A). Principal component analysis and hierarchical clustering confirmed that these two conditions exhibited transcriptome profiles distinct from the remaining CMRL-AT formulations (Fig. 3A and Fig. S6B). Apart from the mannose condition, all CMRL-AT cultures displayed broadly similar transcriptional profiles, with glucose and fructose clustering most closely together, followed by trehalose and the six-sugar condition (C6S).

**Figure 3.**
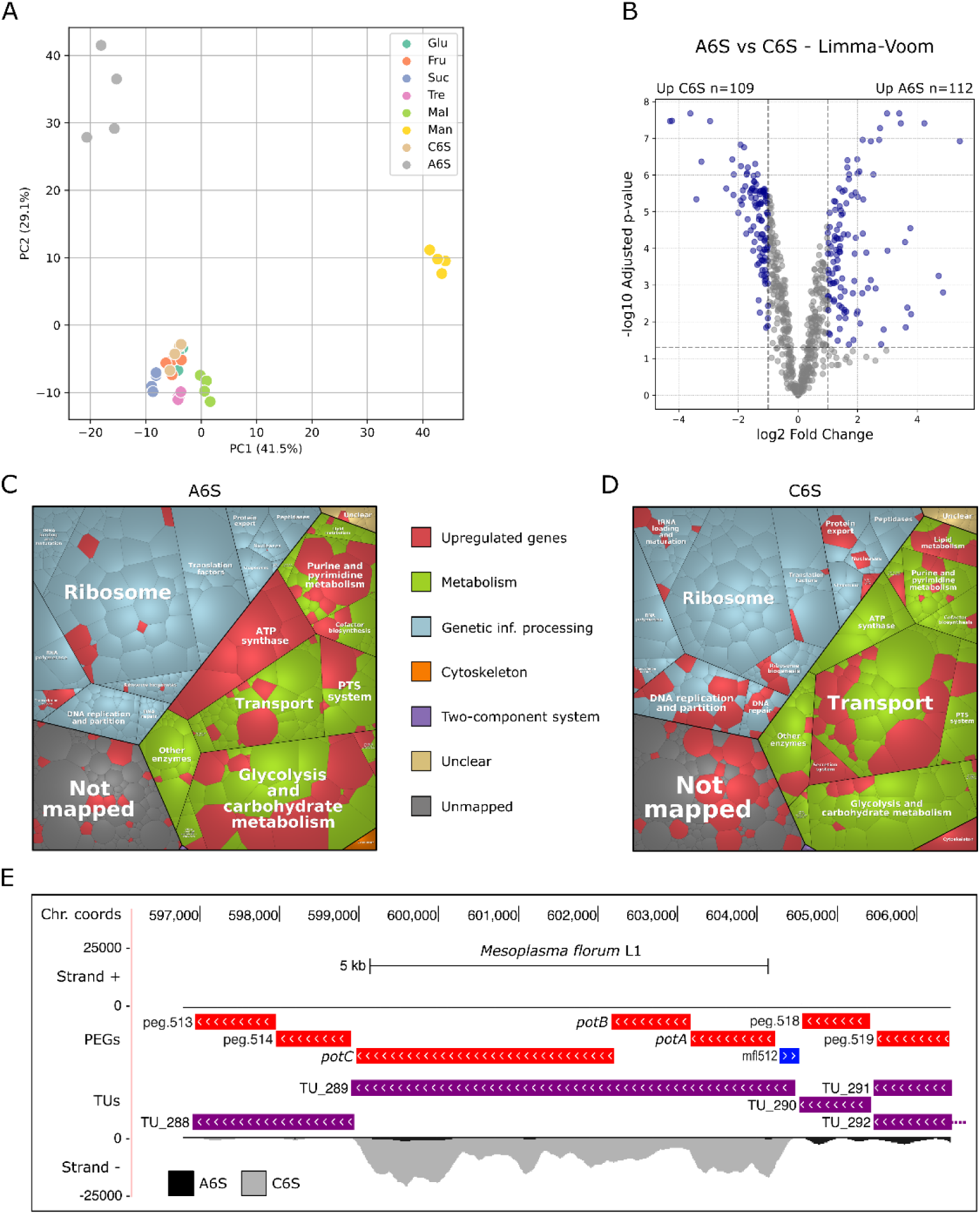
Comparative transcriptome analysis of *M. florum* between complex and serum-free growth media. **A)** Principal component analysis (PCA) performed on all RNA-seq samples analyzed in this study. Glu, CMRL-AT glucose; Fru, CMRL-AT fructose; Suc, CMRL-AT sucrose; Tre, CMRL-AT trehalose; Mal, CMRL-AT maltose; Man, CMRL-AT mannose; C6S, CMRL-AT six sugars; A6S, ATCC 1161 six sugars. **B)** Volcano plot showing differentially expressed PEGs between A6S and C6S conditions, determined by the Limma-Voom package (Law *et al*, 2014). PEGs passing fold change and *p-*value thresholds (gray dotted lines) were considered significantly differentially expressed and are colored in blue. See Materials and Methods for additional details. **C-D)** Voronoi diagrams illustrating the normalized transcription level (median TPM) of *M. florum* PEGs in A6S (C) and C6S (D) growth media. Each polygon represents a specific PEG, area-weighted by its normalized expression. Genes were regrouped into different functional categories updated from a previously published functional hierarchy (Matteau *et al*, 2020) based on the KEGG Orthology (KO) database (Kanehisa *et al*, 2016). Upregulated PEGs in each growth medium are colored in red. **E)** Example of an *M. florum* genomic locus showing differential expression of co-transcribed genes in A6S versus C6S (*potABC* operon). Median strand-specific RNA-seq signal is shown for both conditions (A6S, black; C6S, gray), smoothed over a 16-pixel window. Enhanced RAST annotation is shown, along with previously published *M. florum* transcription units (TU) (Matteau *et al*, 2020). Colored dotted lines indicate genes cropped for representation purposes.

### Transition from complex to serum-free medium induces extensive transcriptome remodeling

To investigate the transcriptional adaptation of *M. florum* to serum-free growth, differential-expression analysis was performed between A6S and C6S. A total of 221 PEGs met the selected significance criteria (|log2 fold change| > 1 and adjusted p < 0.05), corresponding to approximately one-third of all annotated genes (Fig. 3B and Table S4). Of these, 109 PEGs showed higher expression in C6S and 112 in A6S, indicating extensive transcriptome remodeling between the two growth conditions. The pairwise differential expression results were robust to the analytical method, with highly correlated fold-change estimates obtained using Limma-Voom (Law *et al*, 2014) and DESeq2 (Love *et al*, 2014) (Fig. S7 and Table S4).

Visualization of expression-weighted cellular functions revealed substantial differences in transcriptome organization between A6S and C6S (Fig. 3C,D and Fig. S8). Differentially expressed genes were distributed across most major functional categories, including energy metabolism, central carbon metabolism, transport, translation, and biosynthesis (Table S5). Functions associated with growth and energy production were generally more prominent in A6S. For example, all genes encoding ATP synthase subunits (peg.108-115) showed higher expression in A6S, with an average fold difference of 2.48 (Fig. 3C,D, Fig. S8 and Fig. S9A). Several key glycolytic genes also displayed large expression differences in favor of A6S, including those encoding glyceraldehyde-3-phosphate dehydrogenase (peg.583), lactate dehydrogenase (peg.600), and fructose-bisphosphate aldolase (peg.646) (Fig. 3C,D, Fig. 4A, Fig. S8 and Fig. S9B-C). In total, 17 of the 40 PEGs assigned to glycolysis and carbohydrate metabolism were preferentially expressed in A6S, and this category was significantly overrepresented among the differentially expressed genes (Table S5). Genes assigned to the ribosome category followed the same pattern, with seven PEGs enriched in A6S and none in C6S (Fig. 3C,D and Fig. S8).

**Figure 4.**
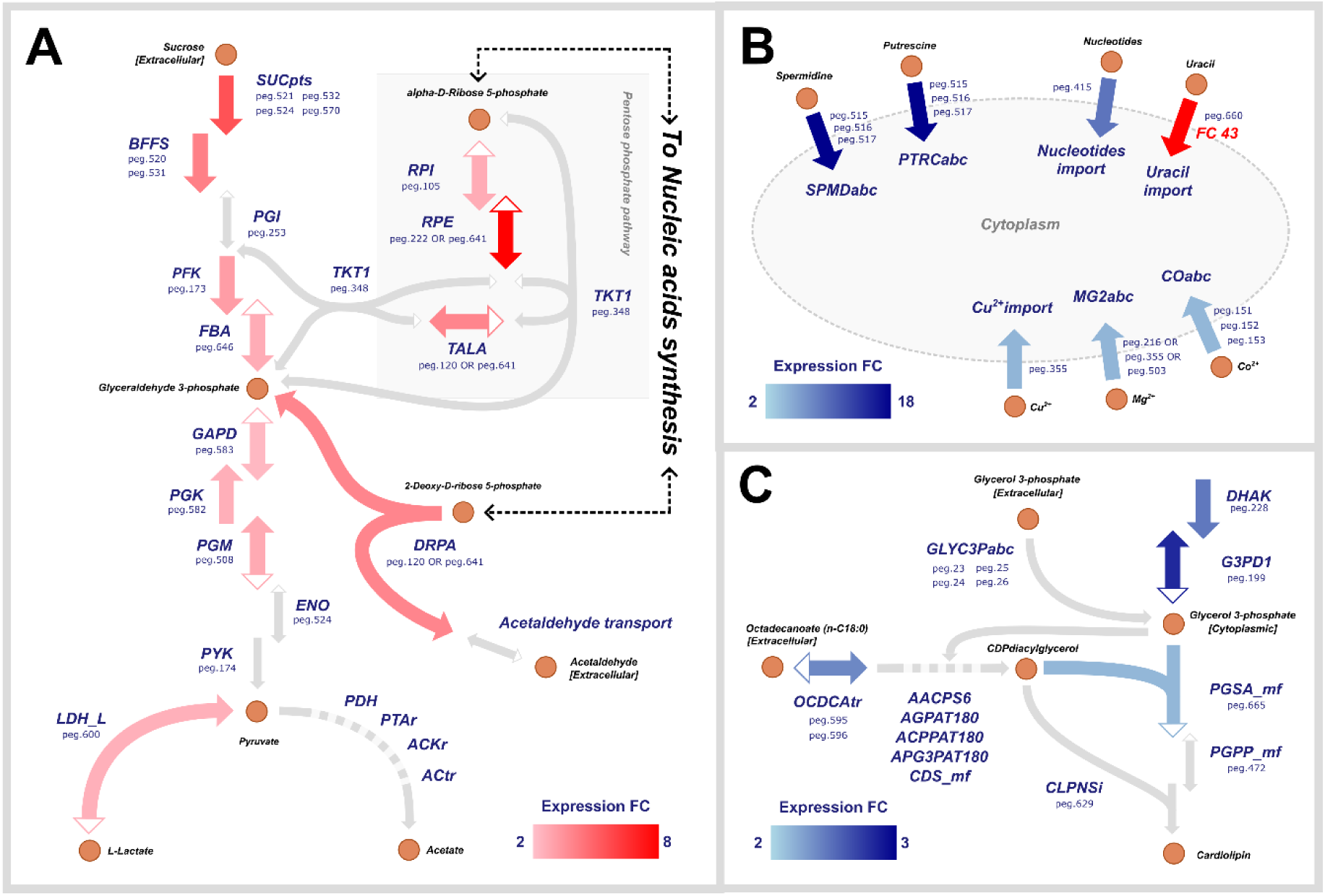
Metabolic map of pathways differentially expressed between CMRL-AT and ATCC. Nodes represent metabolites and arrows represent reactions, pointing in the direction of the reaction. Metabolites are labeled in black; reactions and their associated genes are labeled in dark blue. Reaction color reflects the expression fold-change. **A)** Glycolysis. **B)** Import of critical metabolites. **C)** Lipid degradation.

Distinct trends were observed in C6S, where functions associated with nutrient acquisition were strongly enriched. The transport category contained numerous genes induced in the serum-free medium, most notably the polyamine transporter operon potABC (peg.515-517), whose three components showed the largest expression differences in favor of C6S (Fig. 3D-E, Fig. S8 and Table S4). Transcript levels of potA, potB, and potC were 19.6-, 18.7-, and 12.2-fold greater, respectively, in C6S than in A6S (Table S4). This response extended beyond polyamine transport. Multiple genes associated with cofactor and metal-ion uptake in the iJL208 genome-scale metabolic model (Lachance *et al*, 2021) also showed higher expression in C6S, including transport functions predicted to involve cobalt (peg.151-153), riboflavin (peg.581), magnesium (peg.216, peg.355 and peg.503), and copper (peg.355) (Fig. 3D, Fig. 4B, Fig. S8B, Fig. S9D and Table S4). Consistent with this broader enrichment of transport functions, the nucleobase transporter azgA/pbuG (peg.415) was induced 9.45-fold in C6S, whereas the alternative transporter pyrP (peg.660) displayed one of the largest expression differences in favor of A6S, with a 43.1-fold difference (Fig. S8 and Table S4). Together, these results indicate higher transcription of transport systems involved in the acquisition of polyamines, cofactors, metal ions, and nucleobases during growth in the serum-free medium.

Lipid metabolism was also significantly enriched in C6S, with one-third of the genes assigned to this category (7 of 21 PEGs) meeting the differential-expression criteria, consistent with the expression-weighted Voronoi analysis (Fig. 3D and Fig. S8B). These genes included PEGs predicted to participate in fatty acid transport (peg.595 and peg.596) and phosphatidylglycerophosphate synthesis (peg.665), a step in cardiolipin biosynthesis (Fig. 4C).

### Energy sources elicit distinct transcriptional responses in *M. florum*

To characterize the influence of energy source on *M. florum* gene expression, we compared transcriptomes from cultures grown in CMRL-AT supplemented with glucose, fructose, sucrose, trehalose, maltose, or mannose. Notably, mannose produced a markedly distinct global transcriptional profile (Fig. S6) and was therefore analyzed separately to prevent its dominant response from obscuring differences among the remaining sugar conditions. Pairwise comparison of the mannose and C6S conditions identified 361 differentially expressed PEGs, corresponding to approximately half of the 694 genes in the multi-source annotation. Of these, 170 showed higher expression with mannose, whereas 191 showed higher expression in C6S (Fig. S10A and Table S4). Genes preferentially expressed in C6S were enriched for functions associated with translation and energy metabolism, including ribosomal proteins, translation factors, and ATP synthase subunits. In contrast, genes preferentially expressed under mannose supplementation were enriched for glycolysis and carbohydrate metabolism, with 20 of the 40 genes assigned to this category meeting the differential-expression criteria (Table S6). These results establish mannose as a distinct transcriptional condition among the energy sources tested.

We next examined condition-dependent expression patterns across the five remaining individual sugar conditions. For each gene, we computed a sugar-specificity score (SSS) and clustered the resulting profiles (see Methods). The clustering identified 22 clusters of PEGs with similar expression profiles (Fig. 5A-F, Fig. S11A and Table S8). From them, clusters 1, 2, 3, 4, 6, 11 and 14 displayed more pronounced sugar-associated expression patterns, whereas the six largest clusters (depicted in gray) contained most PEGs with relatively stable expression across conditions (Fig. 5A-F). Pairwise differential-expression analyses were broadly consistent with the cluster-specific patterns (Fig. S10B-E).

**Figure 5.**
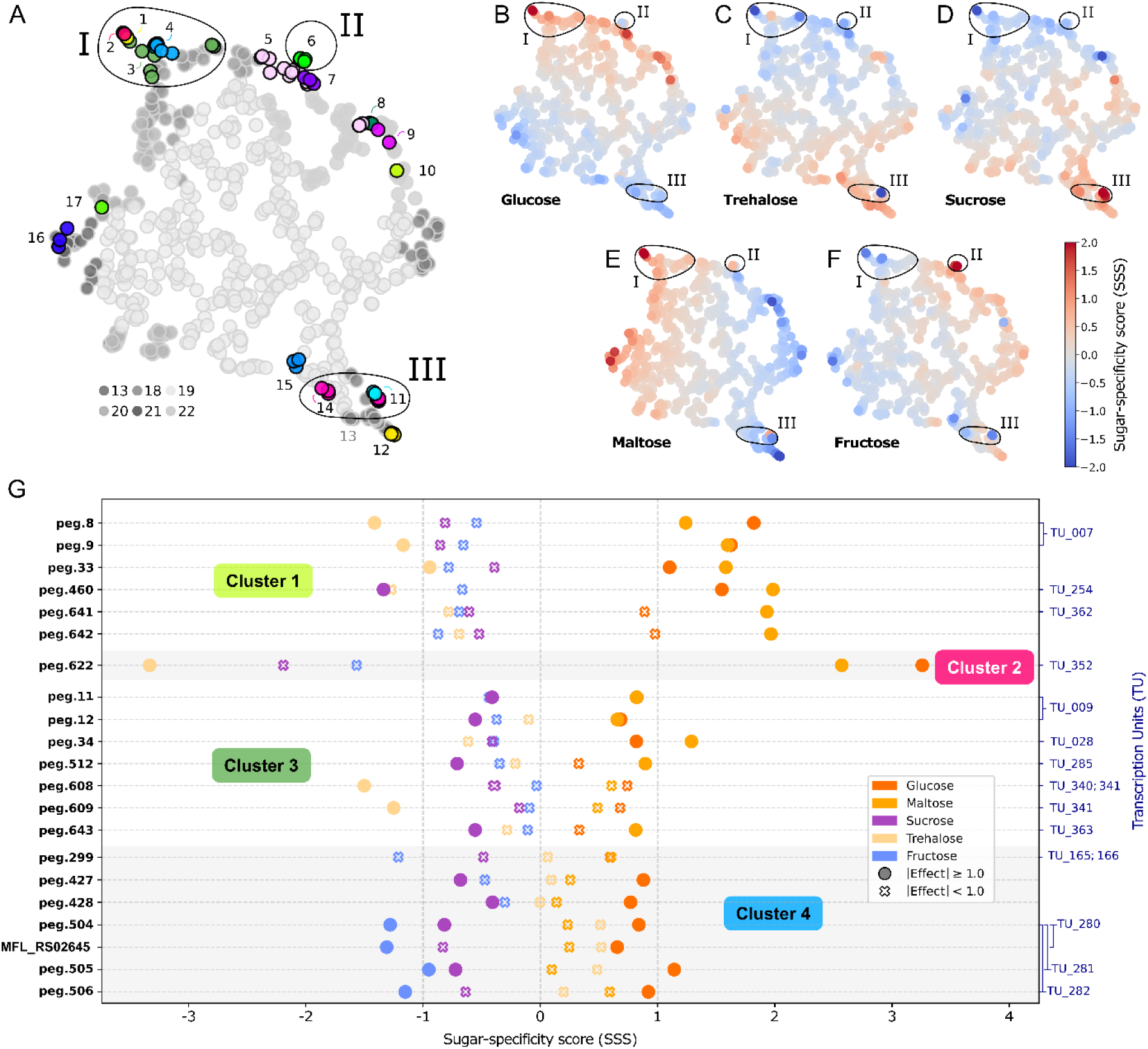
UMAP-based visualization and hierarchical clustering of sugar-specific transcriptional profiles. **A-F)** UMAP embedding of all genes based on their sugar-specific score (SSS) profiles across five sugar conditions (glucose, trehalose, maltose, fructose, sucrose). Each point represents one gene while encircled areas highlight the zones where the groups of interest belong. Group I encompasses Cluster 1-2-3 and 4, Group II includes Cluster 6 and Group III regroups Cluster 11 and 14. **A)** Genes assigned to any cluster with a SSS span ≥ 1.5 are highlighted with distinct colors and labeled by cluster index. Points of different grey scales correspond to genes in low-span clusters of no further analytical interest, representing the baseline from which the clusters of interest are dragged from. **B-F)** Genes coloration represent their SSS profiles across the five sugar conditions: (B) glucose, (C) trehalose, (D) sucrose, (E) maltose and (F) fructose. **G)** Cleveland-style dot plot for Group I, illustrating detailed gene-level SSS values for each sugar condition. Each point represents a gene-condition pair; solid circle markers indicate an absolute effect size ≥ 1.0, while empty cross markers indicate an absolute effect size under 1.0, distinguishing shifts that are large relative to within-condition dispersion from those that are not. Dotted lines at SSS of −1 and 1 are included as references even though there is no formal threshold. Brackets on the right denote transcription unit (TU) membership (Matteau *et al*, 2020)

The clusters in Group I include a total of 21 PEGs and displayed expression profiles generally associated with glucose, maltose, and trehalose supplementation (Fig. 5B-G and Fig. S11B). The strongest response was observed for the single-gene Cluster 2, which displayed strong positive specificity in glucose and maltose, and strong negative specificity in sucrose and trehalose. Cluster 4 included genes predicted to participate in sugar transport and catabolism, as well as the previously unannotated hypothetical gene MFL_RS02645. The expression profile and genomic position of MFL_RS02645 were consistent with its inclusion in the same transcriptional unit (TU) as peg.504-506 (Fig. S12A) previously identified (Matteau *et al*, 2020). Together, these expression patterns identified candidate loci associated with glucose, maltose, and trehalose utilization.

Group II, composed solely of Cluster 6, contains five PEGs displaying a coordinated fructose-associated expression profile (Fig. 5A-F and Fig. S11C,E (top)). All five PEGs showed positive strong specificity under fructose supplementation. Pairwise comparison with glucose further supported this response, with peg.212 and peg.213 displaying log₂ fold changes of approximately −2 in fructose (Fig. S10B). Together, the coordinated expression and predicted functions of these genes support the assignment of Group II as a candidate fructose-responsive transport and catabolic module.

Group III, comprising Clusters 11 and 14 displayed expression profiles mainly associated with sucrose supplementation (Fig. 5A-F and Fig. S11D,E (bottom)). Cluster 11 comprised two genes, peg.531 and peg.532. Both genes exhibited a pronounced strong positive sucrose-associated and negative trehalose response. Pairwise comparison with glucose supported this pattern, with peg.531 and peg.532 displaying log₂ fold changes of approximately −4 in sucrose (Fig. S10E and Table S4). Their coordinated expression and genomic organization were consistent with their inclusion in the same TU_298 (Fig. S11D,E and Fig. S12C). The seven PEGs assigned to Cluster 14 showed strong positive SSS in sucrose. Together, these patterns identify Cluster 11 as a candidate sucrose-responsive transport and catabolic module, whereas Cluster 14 displayed a response that extended to trehalose.

## Discussion

### CMRL-AT enables controlled physiological studies of *M. florum*

The development of CMRL-AT substantially expands the range of physiological experiments that can be conducted with *M. florum*. Despite the elimination of serum and yeast extract, CMRL-AT supported rapid growth and final cell densities comparable to those achieved in the complex ATCC 1161 medium. It also markedly outperformed the previously developed CSY formulation, reducing the doubling time and raising maximal biomass by approximately one order of magnitude (Fig. 1, Fig. S3 and S4). These improvements are particularly important for experimental studies of a near-minimal bacterium, whose extensive biosynthetic deficiencies create a strong dependence on nutrients supplied by the extracellular environment.

Beyond supporting robust growth, CMRL-AT provides substantially greater control over the nutritional environment than ATCC 1161. Growth in CMRL-AT was strongly dependent on the addition of a metabolizable sugar: in its absence, biomass accumulation remained minimal, whereas ATCC 1161 supported similar biomass accumulation with or without sugar supplementation (Fig. 1, Fig. S3 and S4). The undefined biological ingredients of ATCC 1161 therefore appear to provide sufficient metabolizable carbon to mask the contribution of experimentally added sugars. By contrast, the low background growth observed in CMRL-AT enabled the effects of individual energy sources to be examined directly, making this medium well suited for controlled physiological, transcriptomic, and metabolic studies.

CMRL-AT remains a semi-defined rather than fully chemically defined medium because tryptone is required to sustain robust growth. Nevertheless, replacing horse serum, yeast extract, and heart infusion broth with a simpler formulation containing CMRL-1066, Albumax II, and tryptone substantially reduces medium complexity and improves experimental control. A fully defined formulation could ultimately be generated by replacing tryptone with a defined mixture of amino acids, short peptides, and micronutrients, although identifying the components required to reproduce its growth-promoting activity will require systematic deconvolution. CMRL-AT should therefore be viewed not simply as an alternative cultivation medium, but as an enabling platform for investigating how nutrient availability shapes the physiology and functional organization of *M. florum*.

### Medium composition reshapes growth-associated and membrane-bioenergetic programs

The extensive transcriptional differences between A6S and C6S likely reflect their distinct nutrient compositions and the physiological states they support. *M. florum* grew faster in A6S, and this condition was associated with higher expression of genes involved in glycolysis, ribosomal functions, and nucleotide metabolism (Fig. 1B, Fig. 3B-C and Table S5). The relationship between ribosome allocation and growth rate is well established in bacteria, where faster growth requires greater investment in translational capacity to meet higher protein synthesis demands (Bosdriesz *et al*, 2015). Because glycolysis and fermentation constitute the principal routes of energy production in *M. florum*, the coordinated rise in glycolytic gene expression in A6S is likewise consistent with the greater energetic and biosynthetic demands of rapid growth. Together, these patterns suggest that the complex medium supports a transcriptional program favoring energy production, macromolecular synthesis, and cell proliferation.

The coordinated expression of all eight genes encoding the canonical F₁F₀ ATPase provides a potential link between carbon metabolism and membrane bioenergetics (Fig. S9A and File S5). Although F-type ATPases can operate reversibly, their physiological direction depends on the energetic organization of the cell. In respiratory organisms, proton motive force commonly drives ATP synthesis. By contrast, fermentative Mollicutes generate ATP primarily through substrate-level phosphorylation and appear to hydrolyze ATP through the F₁F₀ complex to export protons, thereby supporting membrane potential, intracellular pH regulation, and ion transport. Experiments in *Mycoplasma gallisepticum* showed that glucose-dependent ATP production supports proton motive force and that inhibition of membrane ATPase activity prevents intracellular alkalinization, supporting an ATP-hydrolyzing, proton-extruding orientation of the complex (Béven *et al*, 2012; Zharova *et al*, 2023; Shirvan & Rottem, 1993). The greater expression of the *M. florum* F₁F₀ ATPase operon in A6S, together with the faster acidification and shorter stationary phase observed in this medium, is therefore consistent with higher ATP-dependent proton extrusion to maintain intracellular pH homeostasis (Fig. 3C-D, Fig. S3A, Fig. S9A and Table S4). In this model, higher glycolytic and fermentative activity increases both ATP availability and acid production, thereby raising demand for ATPase-mediated proton export.

The higher expression of ribonucleotide reductase genes in A6S may similarly reflect the greater biosynthetic demands associated with rapid growth. Ribonucleotide reductase supplies the deoxyribonucleotides required for DNA replication by reducing ribonucleotide precursors, and elevated expression of the *nrdAIB* operon would be consistent with increased demand for DNA synthesis (Fig. 3C-D, Fig. S8, Fig. S9E, and Table S4) (Srinivasan *et al*, 2018). Medium composition may provide an additional, non-exclusive explanation. More than 99% of the nucleotide species supplied by the chemically defined CMRL-1066 component of C6S are present as deoxynucleotides, potentially reducing the need for ribonucleotide reduction. In contrast, nucleotide precursors in A6S are supplied largely through yeast extract, whose nucleic-acid-derived material is expected to be enriched in ribonucleotides relative to deoxyribonucleotides (Nordström *et al*, 1966; Feijó Delgado *et al*, 2013). Greater dependence on ribonucleotide conversion in A6S could therefore contribute to the higher expression of *nrdAIB*. Overall, the coordinated enrichment of glycolytic, ribosomal, ATPase, and nucleotide-biosynthesis functions suggests that A6S supports a growth-associated physiological program integrating carbon catabolism, macromolecular biosynthesis, and intracellular pH homeostasis.

### Differences in medium composition reshape nutrient-acquisition functions

The transition from A6S to C6S was accompanied by higher expression of multiple transport systems, suggesting that growth in CMRL-AT places greater emphasis on nutrient acquisition (Fig. 4B, Table S4 and Table S5). The most pronounced response involved the *potABC* operon, whose three genes showed the largest expression differences in favor of C6S (Fig. 3E). These genes are consistently annotated as components of an ABC transporter for spermidine and putrescine, two polyamines with diverse roles in bacterial physiology (File S5). Because *M. florum* lacks a complete pathway for polyamine biosynthesis, it is expected to obtain these compounds from its environment. Their abundance and bioavailability likely differ substantially between the complex ingredients of ATCC 1161 and the simpler CMRL-AT formulation, potentially explaining the strong transcriptional response of *potABC*. Higher expression of this operon is therefore consistent with enhanced polyamine acquisition in C6S.

This response extended to transport systems predicted to mediate the uptake of metal ions, cofactors, and nucleobases. Genes associated with cobalt, magnesium, copper, and riboflavin transport in the *i*JL208 metabolic model showed higher expression in C6S, suggesting that the availability or chemical context of several micronutrients differs between the two media (Table S4). The C6S-enriched nucleobase transporter *azgA/pbuG* may similarly support the acquisition of nucleotide precursors provided by CMRL-1066. By contrast, expression of the predicted uracil transporter *pyrP* was strongly favored in A6S, consistent with the near-absence of ribonucleoside species in the chemically defined component of C6S (Fig. S8 and Table S4) (Feijó Delgado *et al*, 2013; Nordström *et al*, 1966). These opposing transporter responses may reflect differences in the abundance and identity of extracellular nucleobases rather than a uniform increase in nucleotide uptake under either condition.

Together, these observations support a broader nutrient-acquisition response in CMRL-AT. Differences in the concentrations, chemical forms, or bioavailability of polyamines, trace elements, cofactors, and nucleotide precursors between CMRL-AT and ATCC 1161 may favor higher expression of the corresponding transport systems. Transporter promiscuity and uncertainty in some substrate annotations may also contribute to the observed patterns, particularly for metal-ion transporters. Nevertheless, the coordinated response across several transporter classes indicates that medium composition strongly influences how *M. florum* acquires essential metabolites from its environment. The transporters identified here provide a starting point for deconvoluting the growth-promoting activity of tryptone. Targeted supplementation of CMRL-AT with polyamines, trace metals, and nucleobase precursors, combined with monitoring of the corresponding transporter transcripts, would test whether these compounds account for the observed responses and identify which must be included in a fully defined formulation.

### Sugar-responsive transcription refines PTS substrate assignments

Among the six energy sources tested, mannose produced a transcriptional response divergent enough to require separate analysis, with nearly half the annotated genome differentially expressed relative to the six-sugar condition. The direction of that response is informative: translation and energy-metabolism functions were downregulated while glycolytic and catabolic functions were upregulated, even though *M. florum* reaches comparable cell densities on mannose. Growth on mannose also proceeds without the strong acidification seen with glucose or fructose, as does growth on maltose, indicating reduced fermentative flux and, under our model, a correspondingly lower demand for ATP-dependent proton extrusion. Mannose therefore appears to be a poor energy source, possibly catabolized through promiscuous routes rather than a dedicated pathway (Lachance *et al*, 2021), with the cell investing more transcriptional effort to extract less. That a substrate supporting normal growth can impose such a different metabolic demand shows how much the choice of energy source shapes the transcriptional program, and it is precisely this sensitivity that makes sugar-dependent expression informative for functional annotation.

The controlled manipulation of energy sources in CMRL-AT provided an opportunity to refine the functional organization of the *M. florum* phosphotransferase system (PTS). Existing annotations assign conflicting or incomplete substrate specificities to several PTS components, complicating their integration into the *i*JL208 metabolic model (Lachance *et al*, 2021). By combining sugar-dependent expression patterns with predicted protein domains, genomic organization, previously reconstructed transcription units (Matteau *et al*, 2020), and metabolic context, we assigned several PTS components and associated catabolic enzymes to specific sugar-utilization pathways. The resulting assignments are summarized in Table 1 and incorporated into the proposed PTS model shown in Fig. 6.

**Figure 6.**
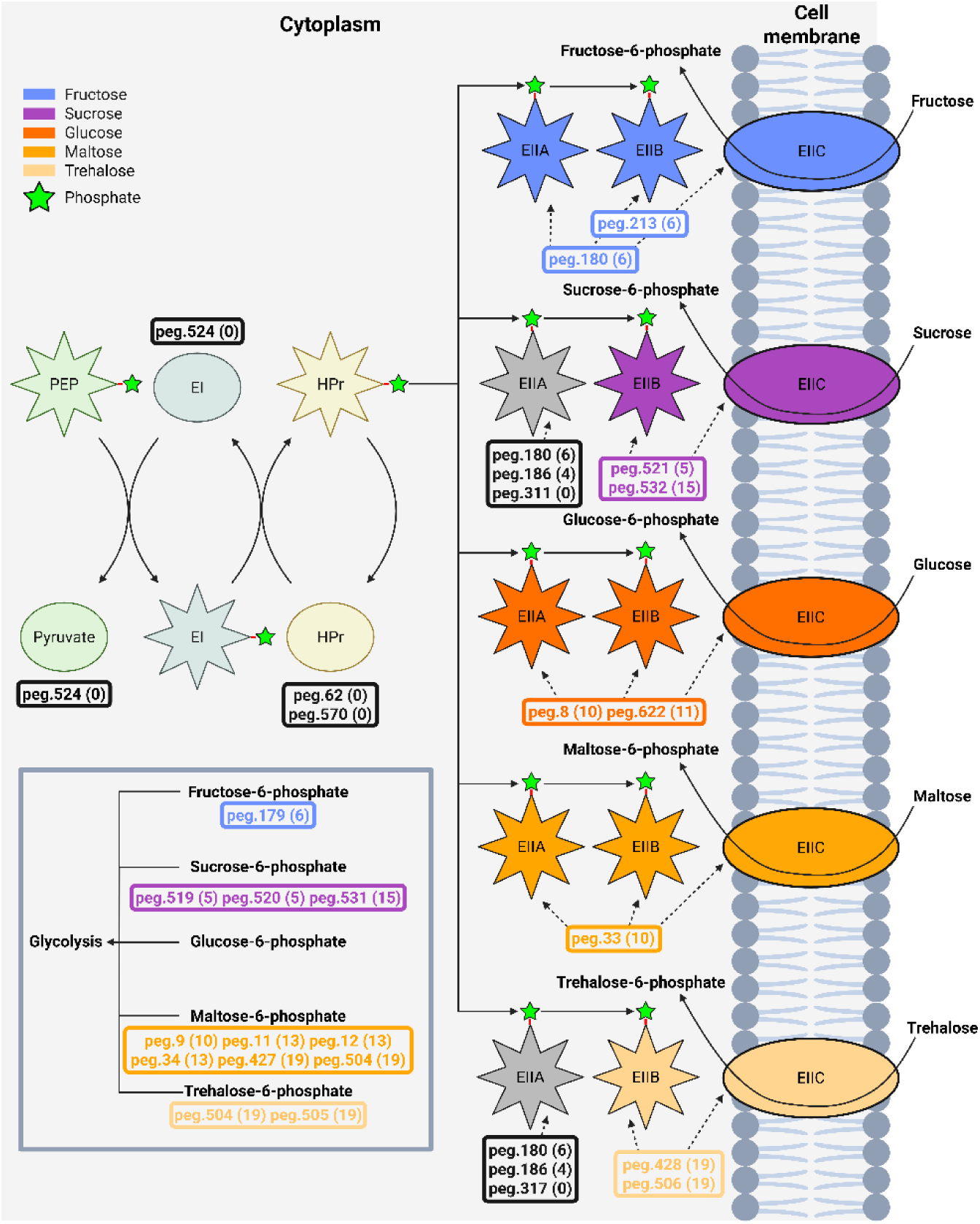
Suggested gene role assignment in the phosphotransferase system (PTS) across five energy sources. A star-shaped protein is phosphorylated, with the small green star that represents the phosphate. The five energy sources retain their color, persistent across this work: glucose (orange), maltose (light orange), sucrose (purple), trehalose (beige), and fructose (blue). On the left-hand side, which represents the cytoplasm, the PTS starts with a non sugar specific protein, phosphoenolpyruvate (PEP) that transfers the phosphate to the enzyme I (EI). The EI transfers the phosphate to an histidine-containing phosphocarrier protein (HPr), which transfers it to the different EIIABC complexes. The corresponding gene(s) are annotated beside the proteins. Annotations in black are non sugar-specific, while colored ones are sugar-specific. The associated cluster is shown beside the PEG name. On the right-hand side, which represents the outside of the cell, are the different available energy sources. With the specific EIIC, the nutrients enter the cell (cytoplasm) while being phosphorylated. The genes processing the different phosphorylated energy sources prior to their entry into glycolysis are depicted in the black box on the bottom left side.

**Table 1.** Revised substrate assignments for *M. florum* phosphotransferase system components. Sugar-responsive transcriptional data were integrated with protein-domain predictions, transcription-unit organization, and metabolic context to refine the annotation of PTS components and associated catabolic enzymes. For each gene, the table reports the proposed domain architecture and preferred substrate alongside the corresponding current annotation. Conclusions are classified as *confirmed* (existing annotation supported by transcriptional evidence), *refined* (substrate range narrowed or specified), *rectified* (previous substrate assignment contradicted by expression data), or *control* (gene used as a reference for assignment validation). Sugar abbreviations: ASC, ascorbate; BGL, β-glucoside; FRU, fructose; GLC, glucose; GlcNAc, *N*-acetylglucosamine; MAL, maltose; SFGLC, glucose superfamily; SUC, sucrose; TRE, trehalose. PTS component nomenclature: EI, enzyme I; HPr, histidine-containing phosphocarrier protein; EIIA, EIIB, EIIC, enzyme II components A, B and C; PEP, phosphoenolpyruvate.

| Gene | Conclusion | Proposed annotation | Preferred sugar | Sugar proposed | Current annotation | Current sugars |
| --- | --- | --- | --- | --- | --- | --- |
| peg.62 | Confirmed | HPr | All | All | HPr | All |
| peg.311 | Refined | EIIABC | SUC | SUC | EIIABC-EIIA-EIIB | SUC, GLC, BGL, SFGLC |
| peg.317 | Refined | EIIABC | MAL | TRE, MAL | EIIABC-EIIA-EIIB | SUC, GLC, BGL, SFGLC |
| peg.524 | Confirmed | PEP, EI | All | All | PEP, EI | All |
| peg.645 | Control | EIIC | ASC | ASC | EIIC | ASC |
| peg.186 | Rectified | EIIA | FRU | SUC, FRU | EIIA | GLC |
| peg.521 | Refined | EIIBC | SUC | SUC, TRE | EIIABC-EIIBC-EIIB | SUC, SFGLC |
| peg.180 | Confirmed | EIIABC | FRU | FRU, MAL | EIIABC | FRU |
| peg.213 | Rectified | EIIBC | FRU | FRU, GLC | EIIABC-EIIBC-EIIC | GlcNAc |
| peg.433 | Refined | EIIBC | MAL | MAL, GLC, FRU | EIIABC-EIIBC-EIIB | SUC, TRE, BGL, SFGLC |
| peg.8 | Refined | EIIABC | GLC | GLC, MAL | EIIABC-EIIA-EIIB | SUC, GLC, BGL, SFGLC |
| peg.33 | Refined | EIIABC | MAL | GLC, MAL | EIIABC-EIIBC-EIIA-EIIB | SUC, GLC, BGL, SFGLC |
| peg.622 | Refined | EIIABC | GLC | GLC, MAL | EIIABC-EIIA-EIIB | SUC, GLC, BGL, SFGLC |
| peg.532 | Refined | EIIBC | SUC | SUC, MAL, GLC | EIIABC-EIIBC-EIIB | TRE, SUC, SFGLC |
| peg.428 | Refined | EIIBC | GLC | GLC, MAL, TRE | EIIABC-EIIBC-EIIB | TRE, SUC, BGL, SFGLC |
| peg.506 | Refined | EIIBC | GLC | GLC, MAL, TRE | EIIABC-EIIBC-EIIB | TRE, SFGLC |
| peg.570 | Confirmed | HPr | All | All | HPr | All |

The most clearly resolved response was associated with fructose. All five genes in Cluster 6 displayed coordinated induction under fructose supplementation. Three of these genes form TU_101, which encodes a predicted DeoR-family transcriptional regulator (peg.178), a phosphofructokinase (peg.179), and a PTS EIIABC component (peg.180). The cluster also contains peg.213, which encodes a predicted EIIBC component but has been variously associated with N-acetylglucosamine or other substrates in existing annotations. Its fructose-associated expression, together with the coordinated response of the other Cluster 6 genes, supports fructose as its physiologically relevant substrate (Fig. 6, Fig. S11C,E-S12B, Table 1 and Table S8-S9). These observations assign Cluster 6 as a fructose-responsive transport and catabolic module and provide a coherent route linking fructose uptake and phosphorylation to glycolysis.

Sucrose supplementation elicited two distinct expression patterns. The selective response of Cluster 11 provides particularly strong evidence for the assignment of peg.531 and peg.532 to sucrose utilization. These genes encode a predicted β-fructofuranosidase and PTS EIIBC component, respectively, and their coordinated induction is consistent with coupled sucrose uptake and cleavage. This transcriptional evidence favors sucrose over the trehalose-related assignments found in some existing annotations (Fig. S11D,E-S12C, Table 1 and Table S8-S9). Cluster 14 also responded to sucrose but showed a less selective pattern that extended to trehalose, suggesting either broader substrate responsiveness or shared regulation between related disaccharide-utilization functions. The data support a dedicated sucrose-associated role for Cluster 11, whereas the functional range of Cluster 14 appears broader and remains less precisely resolved.

The responses associated with glucose, maltose, and trehalose were more complex. Group I displayed overlapping expression patterns under glucose and maltose supplementation, while Cluster 4 additionally responded to trehalose. These clusters contain predicted PTS components and enzymes capable of converting imported sugar phosphates into glycolytic intermediates, including β-glucosidases, glucokinase, and trehalose-6-phosphate hydrolase (Table 1 and Table S8-S9). Their coordinated but partially overlapping responses suggest that *M. florum* combines substrate-preferential transporters with catabolic functions shared among structurally related sugars. In particular, the association of peg.504-506 with Cluster 4 supports a role in disaccharide processing, while the coordinated expression and genomic position of the previously unannotated *MFL_RS02645* strengthen its assignment to the same functional unit. However, the overlapping transcriptional responses do not uniquely resolve every shared EIIA, EIIB, or EIIC component, and some PTS proteins may participate in more than one sugar-utilization pathway (Fig. 6 and Fig. S12A).

Collectively, these assignments reveal a PTS architecture that is more functionally organized, and potentially more flexible, than suggested by any individual annotation source. The strongest substrate associations are supported by convergence among transcriptional selectivity, operon structure, protein-domain predictions, and pathway coherence rather than by expression alone. This integrated evidence establishes well-supported fructose- and sucrose-responsive modules while refining preferential associations for glucose, maltose, and trehalose utilization (Table 1, Fig. 6 and Table S9).

Although direct biochemical transport measurements would further resolve shared or promiscuous components, the revised assignments provide a substantially improved framework for interpreting sugar metabolism in *M. florum* and for updating its genome-scale metabolic reconstruction.

### Environmental context shapes gene function in a near-minimal cell

The sugar-responsive transcriptional programs identified here illustrate an important distinction between genomic minimality and physiological robustness. Most of the PTS-associated genes examined in this study are not considered essential under standard laboratory conditions and are absent from, or frequently removed in, proposed minimal genomes (Fig. 6, Table 1, Table S9 and File S5). Their coordinated responses to different carbohydrates nevertheless indicate that they expand the range of nutritional environments in which *M. florum* can efficiently acquire and metabolize available substrates. Genes that are dispensable in a particular environment may therefore provide substantial adaptive value when nutrient availability changes. In this sense, genome reduction can preserve a core set of functions sufficient for growth under permissive conditions while retaining additional modules that confer metabolic flexibility.

CMRL-AT provides a useful platform for investigating this environmental dependence because its composition can be manipulated without compromising robust growth. By reducing the uncontrolled nutritional inputs present in conventional complex media, it becomes possible to associate specific environmental perturbations with transcriptional, metabolic, and physiological responses. The sugar-dependent expression patterns observed here demonstrate how controlled growth conditions can help resolve gene functions that remain ambiguous from sequence annotation alone. More broadly, these findings emphasize that the functional contribution of a gene cannot be considered independently of the environment in which it is evaluated, particularly when defining minimal genomes or constructing simplified cellular models.

The revised PTS assignments also provide a foundation for improving the *i*JL208 genome-scale metabolic model (Lachance *et al*, 2021) and designing targeted functional studies (Fig. 6, Table 1 and Table S9). Expression-guided pathway assignments can constrain the substrates associated with individual transport systems, clarify how imported sugars enter central metabolism, and identify components that may be shared among multiple pathways. Extending this approach to additional nutrient and stress conditions could further refine the functional annotation of the *M. florum* genome and support condition-specific analyses of gene essentiality. Integrating these data with genetic perturbations and metabolic modeling will ultimately improve our ability to distinguish genes required for basic cellular viability from those that confer robustness across changing environments.

Together, the development of CMRL-AT and the associated transcriptomic analyses establish an experimental framework for studying how a near-minimal bacterium responds to controlled nutritional perturbations. The results reveal broad differences in transcriptional organization between complex and more defined growth environments, identify nutrient-acquisition responses associated with medium composition, and resolve substrate associations within the PTS network. By linking environmental composition to gene expression and functional annotation, this work advances *M. florum* as a tractable model for investigating the organization, flexibility, and design of reduced biological systems.

## Materials and Methods

### Bacterial strains and growth conditions

All experiments were performed using *Mesoplasma florum* strain L1 (ATCC 33453) grown with shaking at a temperature of 34°C in either ATCC 1161, CSY (Lachance *et al*, 2021), or CMRL-AT growth medium. ATCC 1161 is composed of 1.75% (w/v) heart infusion broth, 4% (w/v) sucrose (unless specified otherwise), 20% (v/v) horse serum, 1.35% (w/v) yeast extract, 0.004% (w/v) phenol red and 200 U/ml penicillin G (Matteau *et al*, 2017). CSY medium contains 17.8 g/L of CMRL-1066 chemically defined medium (US Biological, C5900-02A) supplemented with 0.313% (v/v) horse serum, 0.02% (w/v) yeast extract, 6% (w/v) sucrose (unless specified otherwise), and 200 U/ml penicillin G. The optimized version of the CMRL-AT defined medium (see below) is composed of 17.8 g/L CMRL-1066 chemically defined medium (US Biological, C5900-02A), 15 mg/mL Albumax II Lipid-Rich BSA (Gibco, 11021-037), 15 mg/mL Bacto Tryptone (Gibco, 211705), 6% (w/v) sugar (unless specified otherwise), 0.004% (w/v) phenol red, and 200 U/ml penicillin G.

### CMRL-AT medium optimization

To determine the optimal Albumax and tryptone concentrations of the CMRL-AT medium, cell concentrations of three independent *M. florum* cultures prepared in a CMRL-1066 base medium containing 17.8 g/L of CMRL-1066 chemically defined medium (US Biological, C5900-02A), 1% (w/v) sucrose, 0.004% (w/v) phenol red, 200 U/ml penicillin G, and varying concentrations of Albumax II Lipid-Rich BSA (Gibco, 11021-037) and Bacto Tryptone (Gibco, 211705) were measured after 24h and 48h of incubation at 34°C with shaking. Each culture was inoculated with an exponential-phase *M. florum* preculture to obtain an initial concentration of ∼1 × 10^5^ CFU/ml, and cell concentrations were measured by flow cytometry (FCM) as described in (Matteau *et al*, 2020). To determine the optimal sugar concentration, identical experiments were performed but in CMRL-AT medium containing 15 mg/mL Albumax II and 15 mg/mL Bacto Tryptone, but with a varying concentration of either glucose, fructose, sucrose, trehalose, maltose or mannose, or all sugars mixed at equal concentrations.

### Growth kinetics assays

Growth kinetics assays were performed by monitoring the cell concentration of two independent *M. florum* cultures using colony-forming units (CFU) and FCM counts as described in (Matteau *et al*, 2020), except that cultures were incubated for 48h and sampled every hour. Doubling times were evaluated by calculating the growth rate (*r*) between each data point of the exponential portion of the growth curve for each replicate separately. At least three contiguous positive values were considered for each replicate.

Growth rates were calculated according to the following formula:

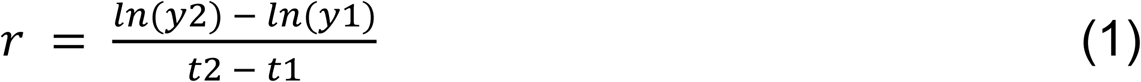

Where *r* is the growth rate, *y*2 and *y*1 the final and the initial cell concentration, and *t*2 and *t*1 the final and initial time, respectively. This equation derives from the equation for exponential growth:

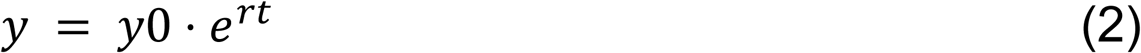

where *y* is the final cell concentration, *y*0 the initial cell concentration, *r* the growth rate, and *t* the time interval. Growth rates were then converted to doubling time according to the following formula:

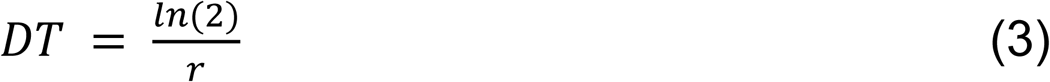

where *DT* is the doubling time and *r* the growth rate.

### RNA-seq experiments, library preparation and sequencing

RNA-seq conditions performed in this study are listed in Table S1. Four biological replicates were performed for each condition. Briefly, for each condition, 20 ml of culture medium was inoculated with an exponential-phase *M. florum* preculture to obtain an initial concentration of ∼1 × 10^5^ CFU/ml. Cultures were incubated at 34°C with shaking for 16 to 20 h to reach the exponential growth phase. Cells were centrifuged for 2 min at 21,000 x *g* and flash frozen in liquid nitrogen until total RNA extraction. Total RNA was extracted by standard hot acid phenol extraction, followed by DNAse I digestion (Zymo Research) at room temperature for 25 min. DNAse-treated RNA was then purified using the RNA Clean & Concentrator-5 kit (Zymo Research) according to the manufacturer’s specifications. RNA concentrations and quality were determined using a 2100 Bioanalyzer instrument (Agilent Technologies). RNA-seq libraries were prepared using the NEBnext Ultra II Directional RNA library kit for Illumina (NEB, 7760) following manufacturer’s specifications with 100 ng of total RNA as starting material for each library. 2 µM of pre-annealed Y-shaped adaptors (Table S7) were used for the adaptor ligation step. RNA-seq libraries were amplified in duplicate by real-time PCR using the NEBnext Ultra II Q5 master mix (NEB, M0544) and the DSN-TruSeq-F / DSN-TruSeq-R primer pair (Table S7) according to the following program: (i) 98°C for 30 sec; (ii) 98°C for 10 sec, 55°C for 30 sec, and 72°C for 20 sec; (iii) repeat previous step until reaching the mid-exponential portion of the amplification curve (∼10-15 cycles); (iv) 72°C for 2 min. Amplification duplicates were pooled and purified using Agencourt AMPure XP beads (Beckman Coulter) with a beads ratio of 0.9. DNA amplicons corresponding to ribosomal RNAs were depleted using the Duplex-specific nuclease as described in (Christodoulou *et al*, 2011). After depletion, libraries were purified again with Agencourt AMPure XP beads (Beckman Coulter) using a beads ratio of 1.6, and then reamplified by real-time PCR using the NEBnext Ultra II Q5 master mix (NEB, M0544) and the MPEX-TruSeq-F / MPEX-TruSeq-R primer pair (Table S7) according to the following program: (i) 98°C for 30 sec; (ii) 98°C for 10 sec, 65°C for 30 sec, and 72°C for 20 sec; (iii) repeat previous step until reaching the mid-exponential portion of the amplification curve (∼15-20 cycles); (iv) 72°C for 2 min. Amplification duplicates were pooled and purified using Agencourt AMPure XP beads (Beckman Coulter) with a beads ratio of 0.9. RNA-seq libraries concentration and quality was finally evaluated on a 5200 Fragment Analyzer system (Agilent Technologies). Paired-end Illumina sequencing (2 x 100 bp) was performed on a Illumina NovaSeq 6000 instrument at the McGill Genome Centre (Montréal, Québec, Canada). Between 4.6 and 54.7 millions of paired-end reads were obtained for each sample, corresponding to an approximate genome coverage of 1,000 to 14,000 (see Table S1).

### RNA-seq data analysis

#### Mesoplasma florum annotation refactoring

To build a comprehensive reference for transcript quantification, four independent genome annotations of *M. florum* L1 were compared: RAST 2018 (Baby *et al*, 2018b), RefSeq 2022 [NC_006055.1], GenBank 2014 [AE017263], and PATRIC 2015 [[ID:265311.5](https://www.bv-brc.org/view/Genome/265311.5)]. Only protein-encoding genes (PEGs) were considered, while rRNAs and tRNAs were excluded because they were depleted during library preparation and are not relevant to the mRNA-focused analysis.

The RAST 2018 annotation served as the reference, and PEGs from the other three sources were compared with it based on their genomic coordinates (start position, end position, and strand). Features with identical start and end coordinates on the same strand were considered matches and merged into a single entry. Features that overlapped a RAST PEG on the same strand but differed at one or both ends were classified as alternative annotations. They were further categorized as N-terminal variants (alternative translation start site), C-terminal variants (alternative stop codon), or N- and C-terminal variants (both ends differ). Features located on the opposite strand or in intergenic regions relative to existing RAST PEGs, which could not be merged with an existing gene model, were added to the annotation as new ORFs. This yielded nine additional PEGs, which were assigned their RefSeq22 MFL-prefixed identifiers when available, otherwise falling back to the GenBank 2014 one (Table S2). All discrepancies were manually curated before the final multi-source annotation was generated (File S5).

For each PEG in the refined annotation, an additive annotation score was computed as the number of sources (1 to 4) in which the feature was identified, where each database had a unique value (RAST = 8, RefSeq = 4, GenBank = 2, and PATRIC = 1). Thus, a unique code was assigned to each combination of annotation sources, allowing the specific sources supporting each PEG to be easily identified (e.g. a gene found in Refseq and GeneBank would have a value of 6 (File S5, column “Annotation_flag”). The agreement between the four databases is depicted in Fig. 2A.

#### Reads processing

Conventional quality assessment steps were performed on raw paired-end reads, including adapter and quality trimming using fastp/0.23.2 (Chen *et al*, 2018). Trimmed reads were aligned to *Mesoplasma florum* L1 reference genome (Baby *et al*, 2013) using Bowtie2/2.3.5 (Langmead *et al*, 2019) and alignment statistics were computed with Samtools flagstat (Danecek *et al*, 2021). Reads with low mapping quality (MAPQ < 10), typically arising from multi-mapping to duplicated loci such as rRNA operons, were removed. Strand-specific genome coverage was computed using Bedtools/2.30 genomecov (Quinlan & Hall, 2010). Gene-level read counts were obtained with featureCounts (Liao *et al*, 2014) using the mentioned refactored genome annotation. The reads processing was boxed in a Snakemake (Mölder *et al*, 2025) pipeline for efficient computing on high-performance computing (HPC) infrastructure.

#### Pairwise differential expression analysis

Pairwise differential expression was assessed using limma-voom (Law *et al*, 2014) and DESeq2 (Love *et al*, 2014) ran with r/4.1.2 (R Core Team, 2021) with r-bundle-bioconductor/3.14, applied independently to the count matrix. For each pairwise comparison, genes with zero counts in any sample were excluded. For limma, library sizes were normalized with the TMM method, and counts were transformed to log2-CPM with voom. Linear models were fitted with lmFit and moderated t-statistics computed with eBayes; p-values were adjusted by the Benjamini–Hochberg method. DESeq2 was run on the same filtered counts with default parameters (median-of-ratios normalization, Wald test, Benjamini–Hochberg adjustment). Significance thresholds for adjusted p-values and absolute fold changes were empirically defined as 0.05 and 2 respectively to determine differentially expressed genes.

Functional enrichment (hypergeometric) analysis was performed using scipy/1.9.3 (Virtanen *et al*, 2020) on functional categories updated from a previously published functional hierarchy (Matteau *et al*, 2020) based on the KEGG Orthology (KO) database (Kanehisa *et al*, 2016).

#### Sugar-specific transcriptional patterns analysis

To characterize sugar-specific transcriptional patterns, a compositional analysis approach was conducted using ALDEx2/1.26.0 (Fernandes *et al*, 2014). Considering that we were focusing on subtle trends, the broadly different mannose condition was excluded from this analysis. To obtain a condition-agnostic baseline, an average expression profile was computed for each gene across all conditions and appended as an additional pseudo-sample. This average served as an anchor, enabling the estimation of expression shifts for each sugar condition relative to the global transcriptional background rather than to a specific condition. Compositional transformation of the count data of each individual condition was performed using ALDEx2’s centered log-ratio (CLR) procedure with 512 Monte-Carlo Dirichlet instances to account for sampling uncertainty. Differential abundance and effect size were computed using aldex.glm.effect (respectively .diff.btw and .effect columns), which fits generalized linear models to CLR-transformed values for each gene across conditions. Effect sizes are expressed as the ratio of between-condition difference to within-condition dispersion. Because the CLR transformation maps compositional data into real space, where Euclidean distances correspond to Aitchison distances, CLR-based values are suitable for distance-based clustering (Fernandes *et al*, 2014). The compositional differential abundance of each condition relative to the average pseudo-sample was aggregated into a matrix and named sugar-specificity score (SSS) for facilitated reading. The SSS matrix captures the magnitude and direction of each gene’s transcriptional shift relative to the global average anchor, while the associated effect size matrix is also consulted to guide interpretation.

#### Dimensionality reduction and clustering of condition-specific expression profiles

To visualize the global structure of condition-specific expression patterns, the SSS matrix was reduced using Uniform Manifold Approximation and Projection (umap-learn/0.5.5; (McInnes *et al*, 2018)) with Euclidean distance, with 10 nearest neighbors, a minimum distance of 0.1, two output components and a fixed random seed (42). To identify groups of genes sharing coherent sugar-specific transcriptional profiles, agglomerative hierarchical clustering was applied to the SSS matrix using sklearn/1.2.0 (Pedregosa *et al*, 2011) with complete linkage and Euclidean distance. The resulting dendrogram was inspected to determine an appropriate partition, and the tree was cut to yield 22 clusters. This number of clusters was set empirically to resolve fine-grained transcriptional patterns among the minority of genes exhibiting meaningful condition-dependent variation, rather than to optimize any formal clustering criterion. As expected, most genes displayed low SSS span and low inter-condition variability, and consequently collapsed into a small number of large, uninformative clusters.

#### Cluster selection and grouping

Clusters in which at least one gene exceeded a SSS span threshold of 1.5 were retained for detailed examination, which means there was likely at least one important difference between two conditions worth mentioning. For each retained cluster, condition-specific transcriptional profiles were visualized using a Cleveland-style dot plot displaying, for each gene, the SSS value for all five sugar conditions along a common axis. This representation allows direct comparison of the magnitude and direction of expression shifts across conditions within and between genes. Each data point was additionally annotated with the ALDEx2 effect size for the corresponding gene-condition pair. Unlike the raw SSS, which reflects only the magnitude of the estimated shift, the effect size simultaneously accounts for within-condition variability: a large SSS accompanied by a low effect size indicates that the observed shift is small relative to the noise inherent to that gene’s expression distribution, and should therefore be interpreted with caution.

Points with an absolute effect size below a threshold of 1.0 were distinguished visually using a different marker, allowing rapid identification of condition-specific signals that are both large in magnitude and robust to within-condition dispersion. Selected clusters were then grouped according to their broad transcriptional patterns, and cluster averages were visualized in radar charts. Along with SSS and effect size, per condition mean normalized counts were inspected to obtain a sense of biological scale involved in each transcriptional shift (Table S8). In regard to all above metrics, we selected the more interesting clusters and bundled them into Groups I to III for further investigation and representation.

#### PTS scheme

The iJL208 metabolic model suggests an approximate function for most genes, but is limited by the depth of references like RefSeq (NC_006055.1) and PATRIC (Genome ID: 265311.5) (Lachance *et al*, 2021). However, the discrepancies between the annotation sources render the model resolution arduous. We queried each of the concerned gene peptidic sequences in protein-specialized databases such as UniProt (The UniProt Consortium *et al*, 2023) and InterPro (Paysan-Lafosse *et al*, 2023) to identify functional domains often unnoticed when relying solely on nucleotide sequence homology. This helps us to address the likely roles of a gene when available gene annotations may be conflicting. In some cases, the annotations are coherent for the function, but lack consensus for the involved sugars. To pinpoint the right preferred sugar for each gene, we integrated our conditional expression data (Fig. 5 & Fig. S11) with the search results from UniProt and InterPro (File S5, Table S8, Table S9).

#### Data visualisation

Metabolic pathway-level visualization was performed using escher/1.7.3 (Buchweitz *et al*, 2020). Read coverage were visualized in a locally hosted UCSC Genome Browser (Casper *et al*, 2026) instance, which is available at http://bioinfo.ccs.usherbrooke.ca/M_florum_transcriptomic_hub.html), facilitated by Python trackhub/0.2.4 module (trackhub, 2017). Other various visualizations were generated using an array of Python packages, including Matplotlib/3.6.2 (Hunter, 2007) and Seaborn/0.12.2 (Waskom, 2021), UpSetPlot/0.8.0 (Lex *et al*, 2014). Further data handling and metric computations were made with in-house Python/3.8 (Python Software Foundation, 2020) scripts, which are available upon request.

## Data availability

### Trackhub

http://bioinfo.ccs.usherbrooke.ca/M_florum_transcriptomic_hub.html

### RNA-seq data

Gene Expression Omnibus GSEXXXXX

## Acknowledgements

The authors would like to thank Frédérique White for contributing initial RNA-seq processing scripts (Limma-Voom and Deseq2). This research project was funded by the Natural Sciences and Engineering Research Council (NSERC) Discovery grant program and the Canada Graduate Research Scholarship – Doctoral program. Access to computational resources was provided in part by Calcul Québec (http://www.calculquebec.ca) and Digital Research Alliance of Canada (https://alliancecan.ca).

## Author contributions

AD: Conceptualization, Formal analysis, Investigation, Methodology, Writing - original draft

JG: Data curation, Formal analysis, Methodology, Visualization, Writing – original draft, Writing - review and editing

SJ: Data curation, Formal analysis, Methodology, Validation, Visualization, Writing - original draft, Writing - review and editing

DM: Conceptualization, Data curation, Formal analysis, Funding acquisition, Investigation, Methodology, Supervision, Validation, Visualization, Writing - original draft, Writing - review and editing

P-ÉJ: Conceptualization, Supervision, Resources, Funding Acquisition, Writing - review and editing

SR: Project administration, Conceptualization, Supervision, Resources, Funding Acquisition, Writing - review and editing

## Expanded View Figure legends

**Figure S1.**
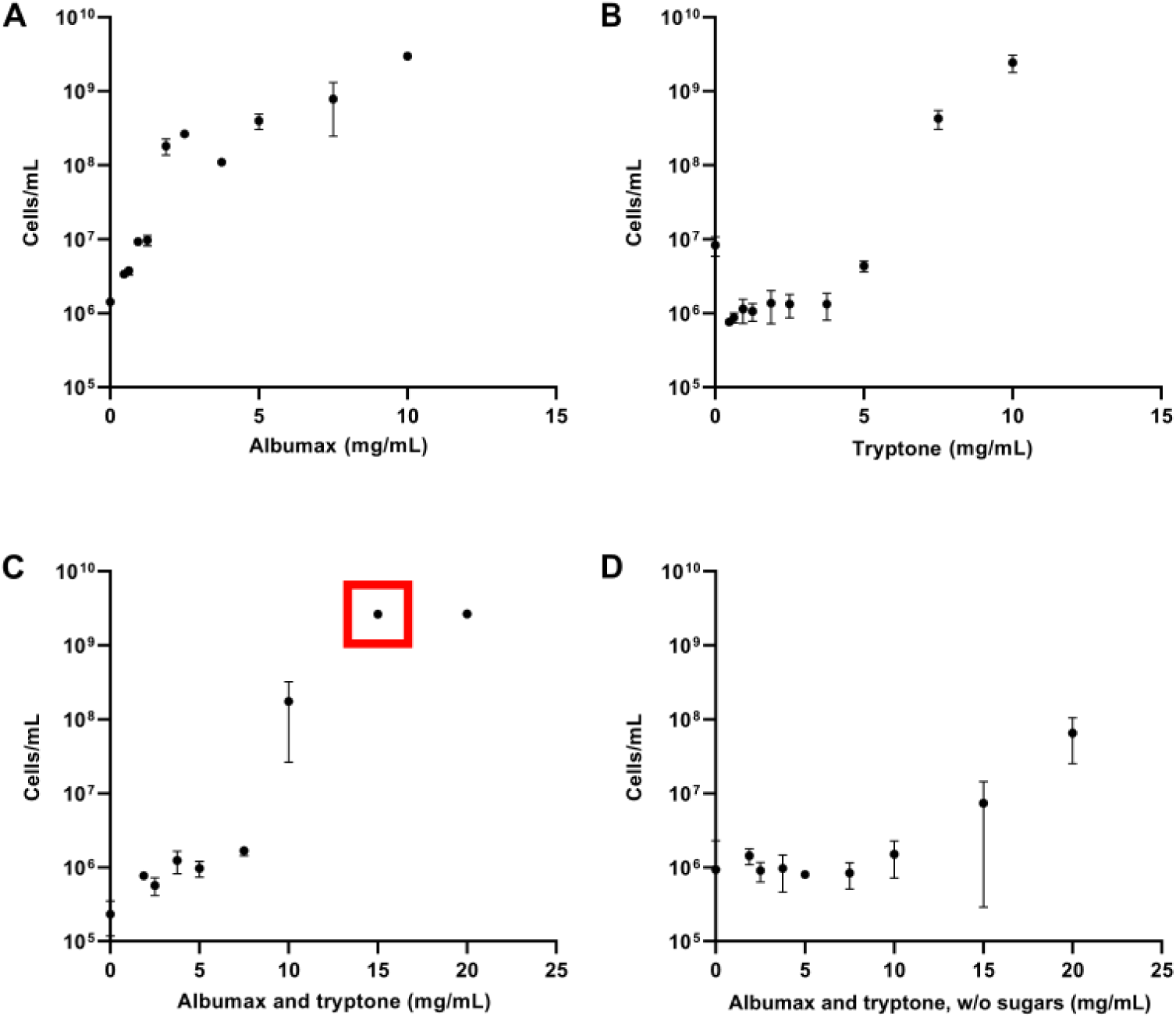
Impact of Albumax and tryptone supplementation on *M. florum* growth in CMRL-AT medium. Cell concentrations were determined by flow cytometry after 48h of incubation. **A-D)** Growth in CMRL-AT supplemented with varying concentrations of Albumax II (A), tryptone (B), both (C), or without sugar (D) to evaluate possible sugar contamination in Albumax and tryptone supplements, red box shows the Albumax II and tryptone concentrations chosen. Data points represent the mean and standard deviation of biological triplicates.

**Figure S2.**
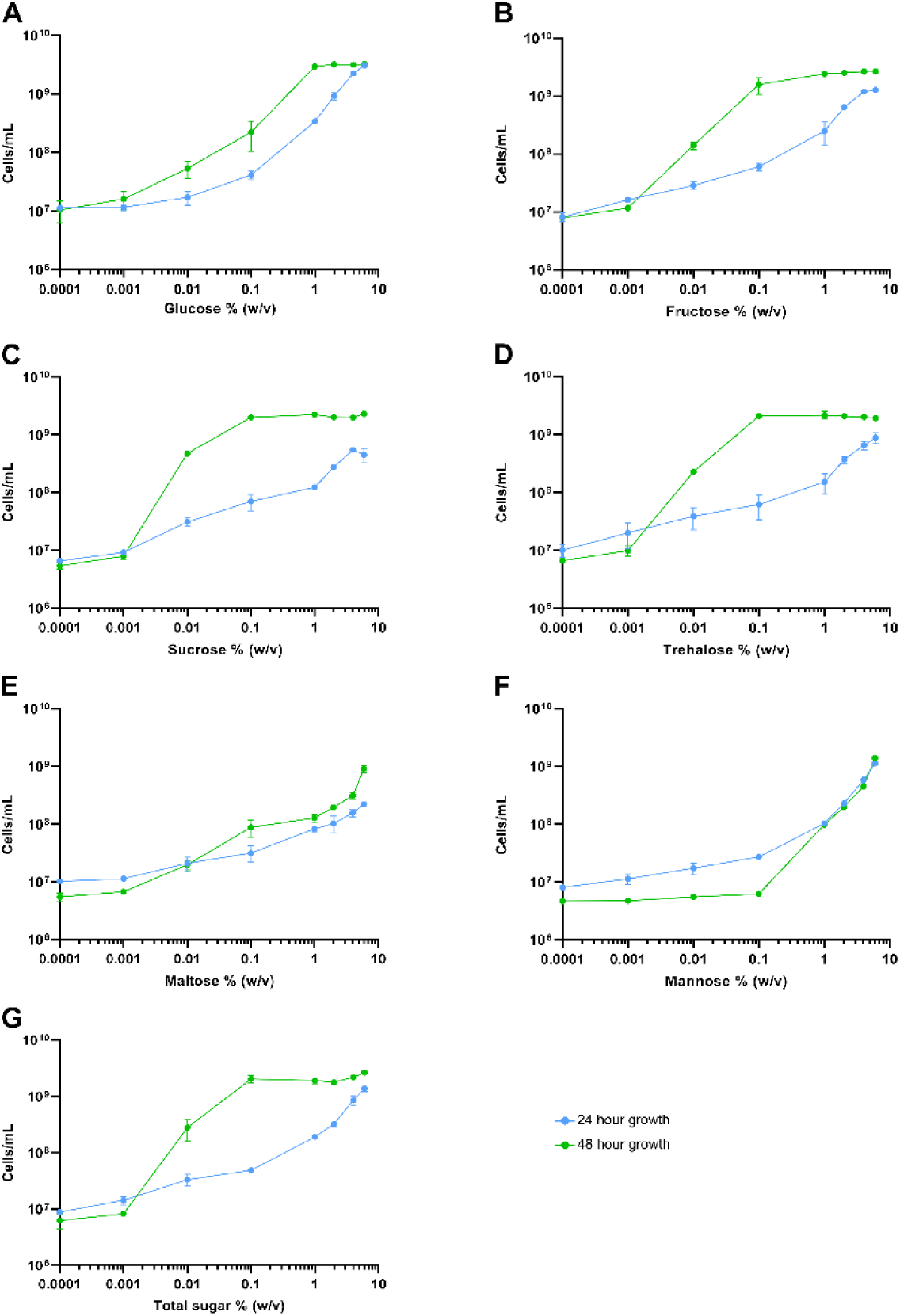
Impact of the concentration of various sugars on *M. florum* growth in CMRL-AT medium. Cell concentrations were determined by flow cytometry after 24 and 48h of incubation in CMRL-AT supplemented with varying concentrations of each sugar. **A)** Glucose. **B)** Fructose. **C)** Sucrose. **D)** Trehalose. **E)** Maltose. **F)** Mannose. **G)** All six sugars combined at equivalent concentrations. Data points represent the mean and standard deviation of technical triplicates.

**Figure S3.**
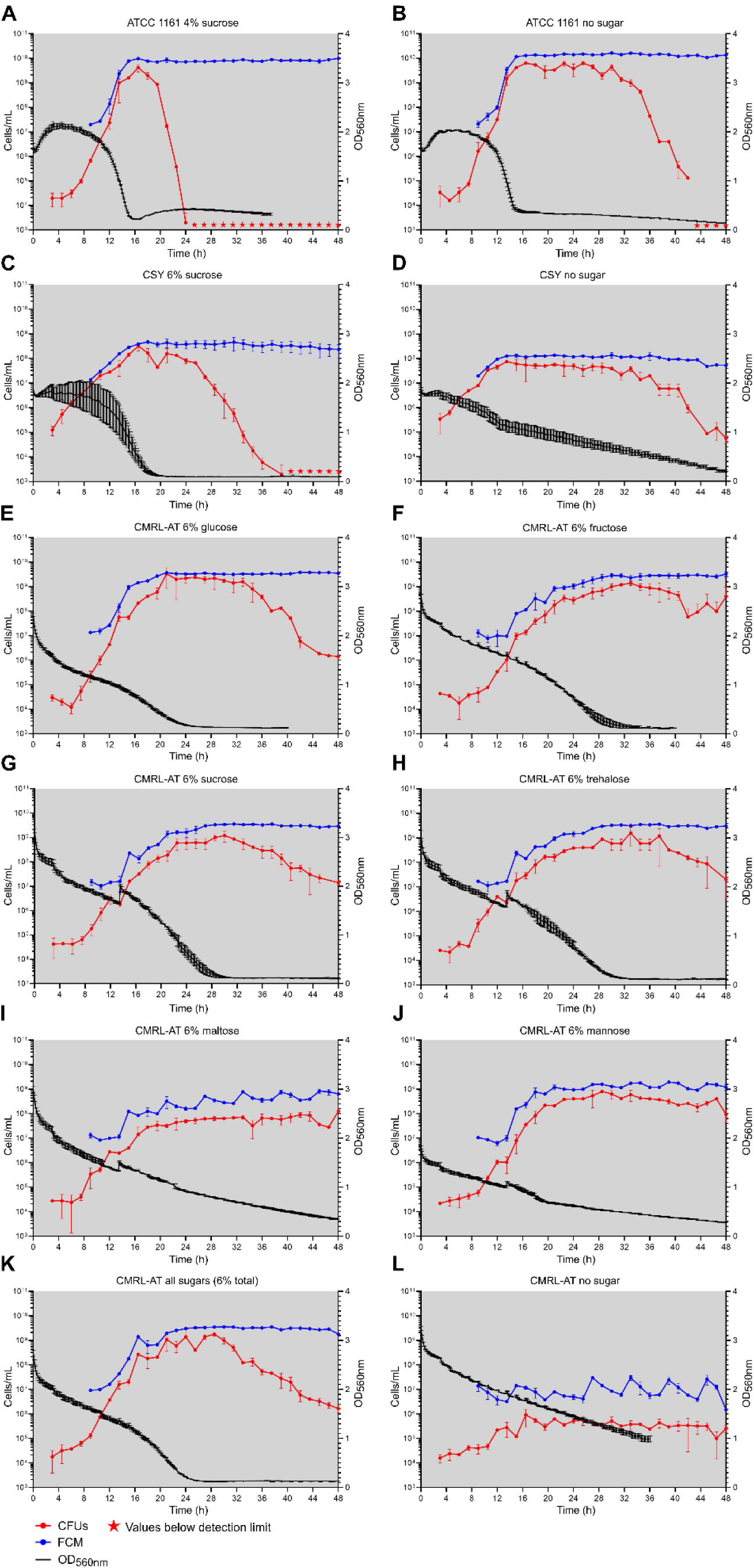
*M. florum* growth kinetics in different growth media. Growth was monitored at 34°C for 48h by measuring optical density at 560 nm (black circles) and cell concentrations using two methods, colony-forming units (CFU, red squares) and flow cytometry (FCM, blue triangles). **A)** ATCC 1161 with 4% sucrose. **B)** ATCC 1161 without sugar. **C)** CSY with 6% sucrose. **D)** CSY without sugar. **E)** CMRL-AT with 6% glucose. **F)** CMRL-AT with 6% fructose. **G)** CMRL-AT with 6% sucrose. **H)** CMRL-AT with 6% trehalose. **I)** CMRL-AT with 6% maltose. **J)** CMRL-AT with 6% mannose. **K)** CMRL-AT with all six sugars at equivalent concentrations. **L)** CMRL-AT without sugar. The dots and error bars indicate the mean and standard deviation values obtained from biological duplicates. Stars indicate CFU values below the detection limit of the experiment.

**Figure S4.**
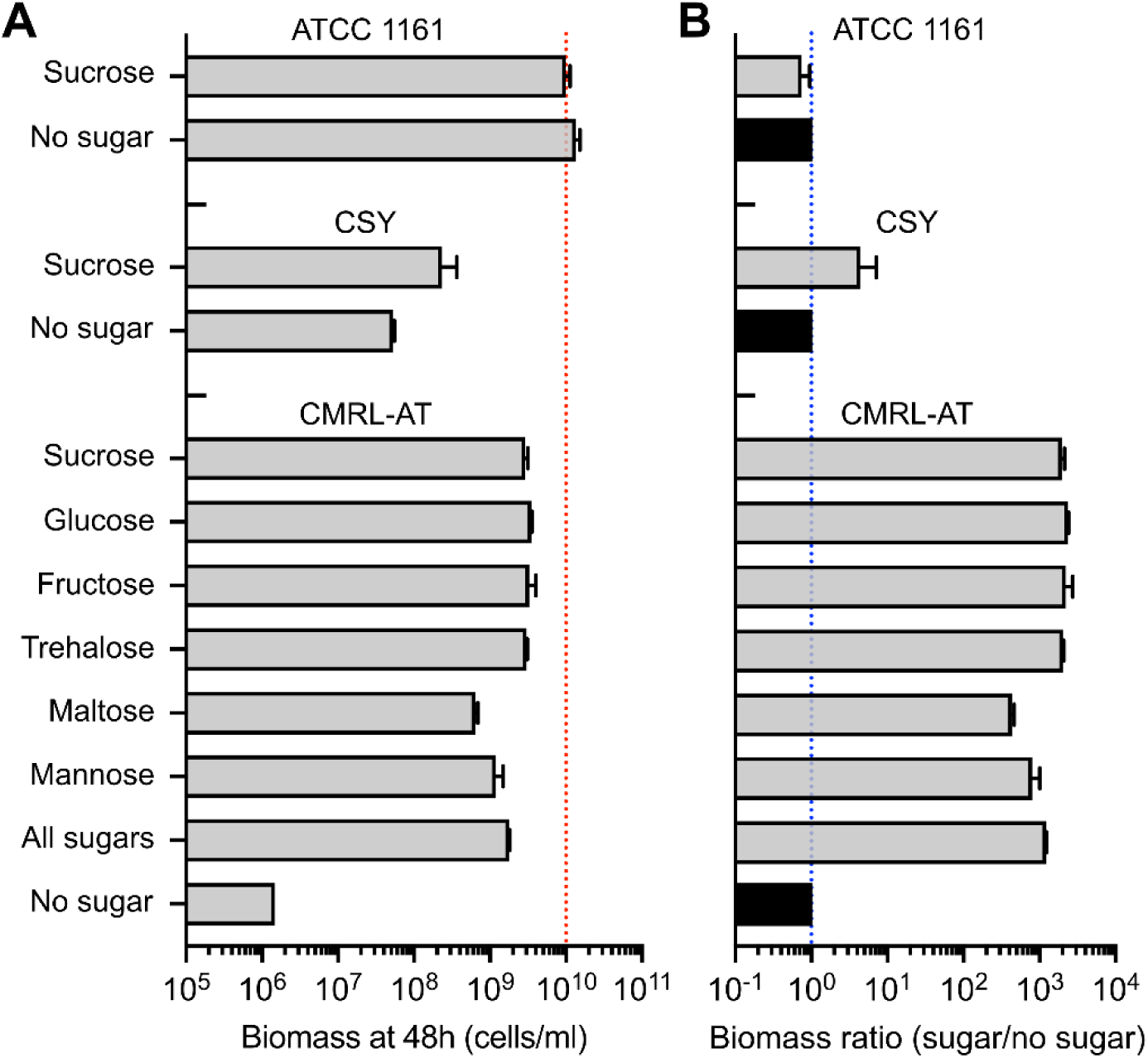
*Mesoplasma florum* biomass after 48h of incubation in different media. **A)** *M. florum* biomass at 48h of incubation in three different media (ATCC 1161, CSY, and CMRL-AT), with or without sugar supplementation, determined by flow cytometry. The red dotted line indicates the value obtained for the ATCC 1161 medium with 4% sucrose. See Figure S3 for complete growth curves. **B)** Biomass ratio at 48h of incubation between sugar supplemented and non-supplemented conditions determined for each growth medium. Growth conditions without sugars were arbitrarily set to a ratio of 1, as also denoted by the blue dotted line. Bars and error bars indicate mean and standard deviation calculated over biological duplicates, respectively.

**Figure S5.**
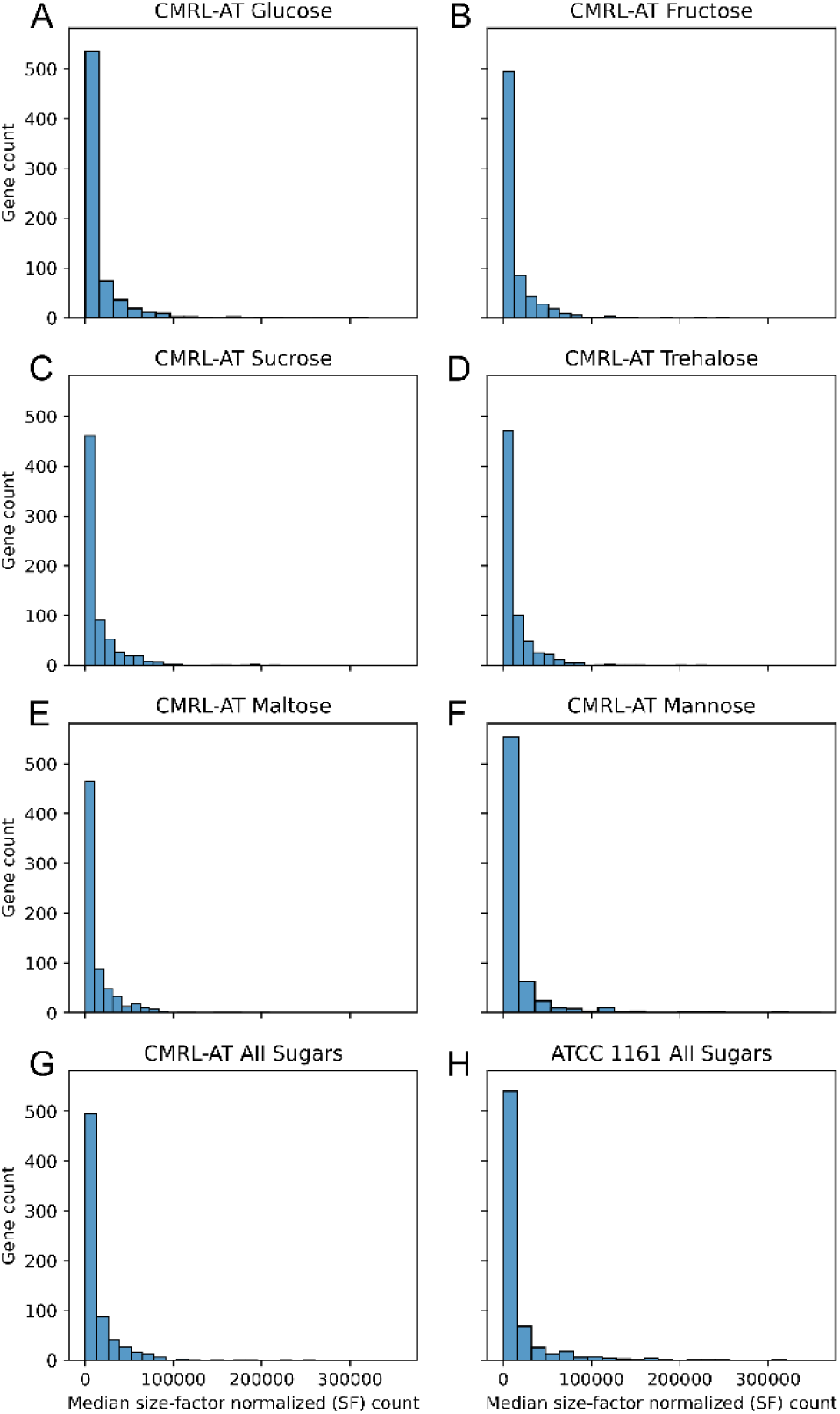
Distribution of gene expression in various conditions. Transcription level of every *M. florum* protein-encoding gene was evaluated by RNA-seq in CMRL-AT supplemented with either 6% glucose, fructose, sucrose, trehalose, or maltose, as well as in CMRL-AT and ATCC 1161 supplemented with 1% of each sugar, for a final concentration of 6%. Median size-factor-normalized (SF) calculated from four biological replicates was used to determine transcription level in each condition.

**Figure S6.**
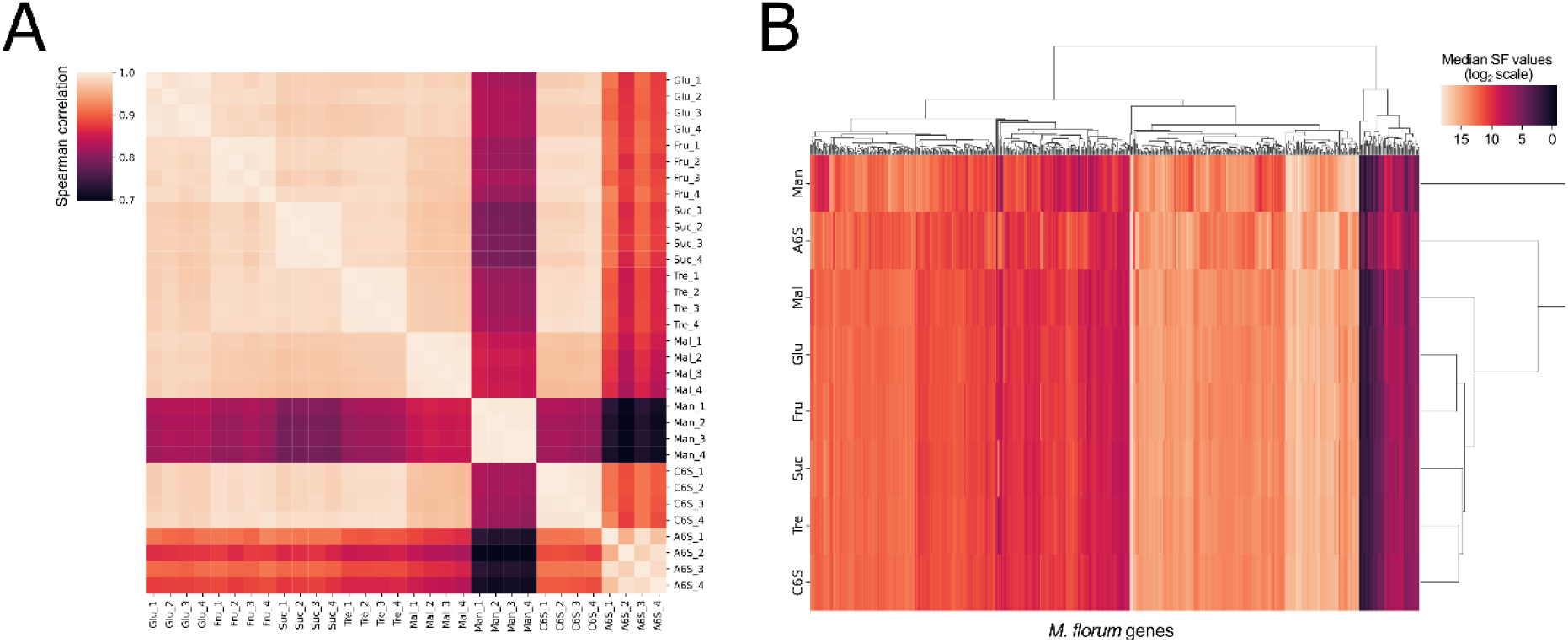
Correlation matrix and principal component analysis of RNA-seq samples. Sample comparisons were performed on size-factor-normalized and scaled counts. **A)** Pearson’s correlation heatmap of all RNA-seq conditions and replicates generated in this study (see Table S1 for the complete list). **B)** Genome-wide transcription level of *M. florum* protein-encoding genes (PEGs) in ATCC 1161 medium and in CMRL-AT with different sugar supplementation. Genes were clustered and colored according to their median size-factor normalized expression count (SF) calculated over 4 replicates. The enhanced RAST annotation developed in this study was used as a reference (see Table S5). Glu, CMRL-AT glucose; Fru, CMRL-AT fructose; Suc, CMRL-AT sucrose; Tre, CMRL-AT trehalose; Mal, CMRL-AT maltose; Man, CMRL-AT mannose; C6S, CMRL-AT six sugars; A6S, ATCC 1161 six sugars.

**Figure S7.**
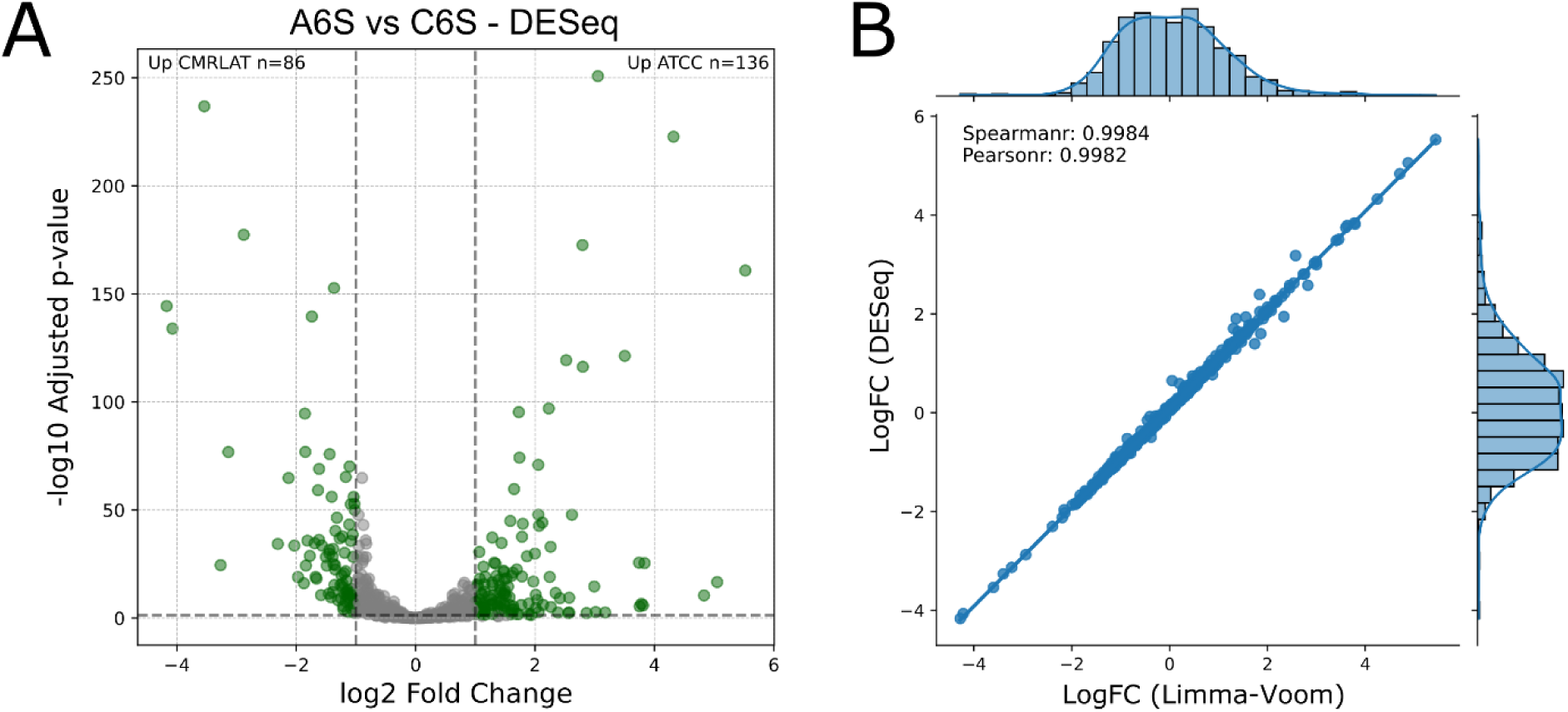
Differential expression analysis using DESeq2. **A)** Volcano plot comparing expression of protein-encoding genes (PEGs) between A6S and C6S conditions, determined by the DESeq2 package (Love *et al*, 2014). PEGs passing both the fold change (|log2FC| ≥ 1) and adjusted *p*-value (BH-adjusted ≤ 0.05) thresholds (gray dashed lines) are considered differentially expressed genes (DEGs) and are colored in green. See Materials and Methods for additional details. **B)** Comparison of log2 fold changes determined by Limma-Voom (Law *et al*, 2014) and DESeq2 (Love *et al*, 2014) packages. Spearman and Pearson correlation coefficients are indicated, along with marginal histograms showing the distribution of fold-change values for each method.

**Figure S8.**
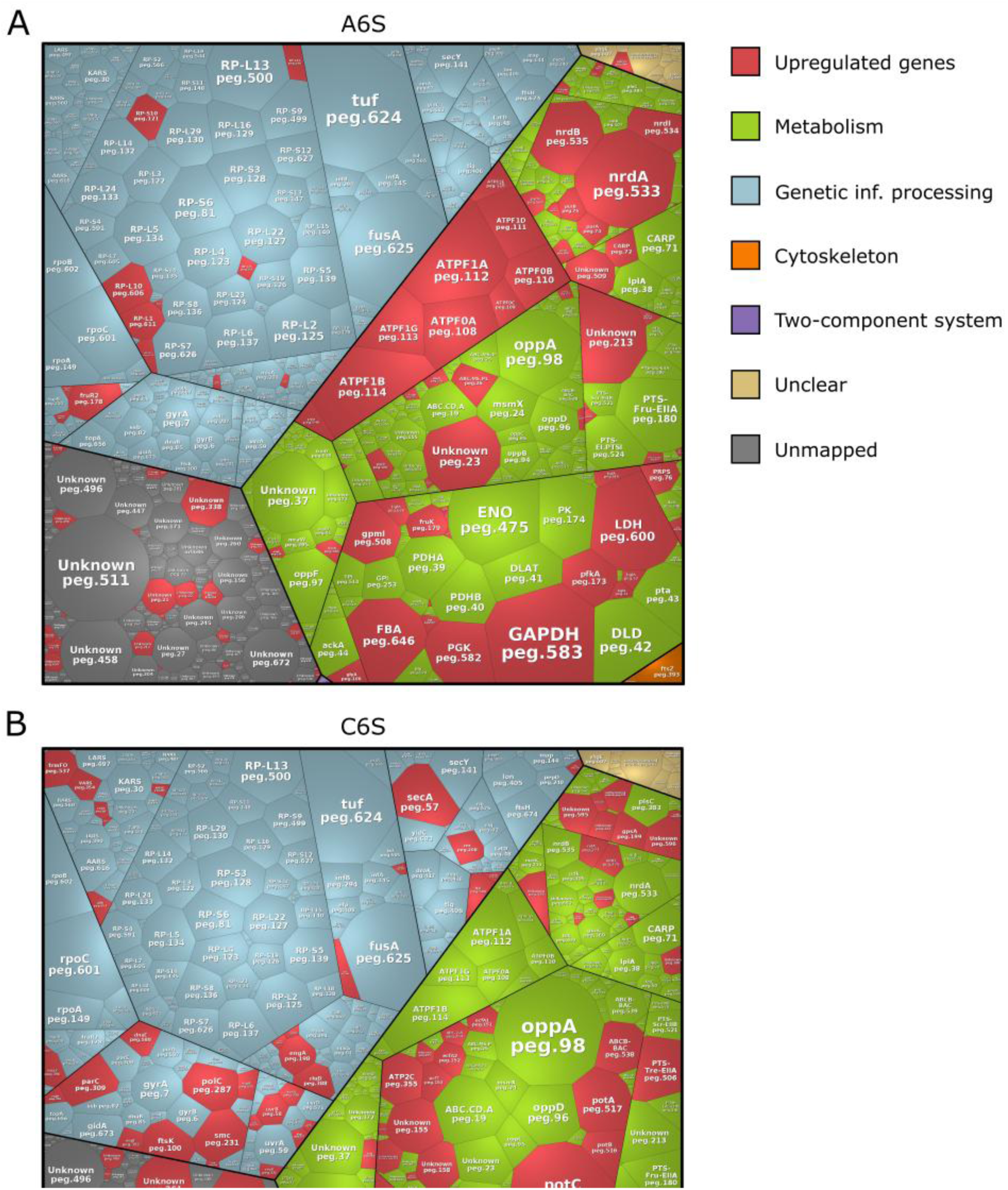
Detailed view of Voronoi diagrams illustrating gene expression in A6S and C6S growth media. Each Voronoi diagram illustrates the normalized transcription level (median TPM) of *M. florum* PEGs. **A)** A6S growth medium. **B)** C6S growth medium. Genes were regrouped into different functional categories updated from a previously published functional hierarchy (Matteau *et al*, 2020) based on the KEGG Orthology (KO) database (Kanehisa *et al*, 2016). Each polygon represents a specific PEG area-weighted by its normalized expression quantified by RNA-seq, identified by its corresponding RAST locus tag and KEGG gene product. Upregulated PEGs in each growth medium are colored in red.

**Figure S9.**
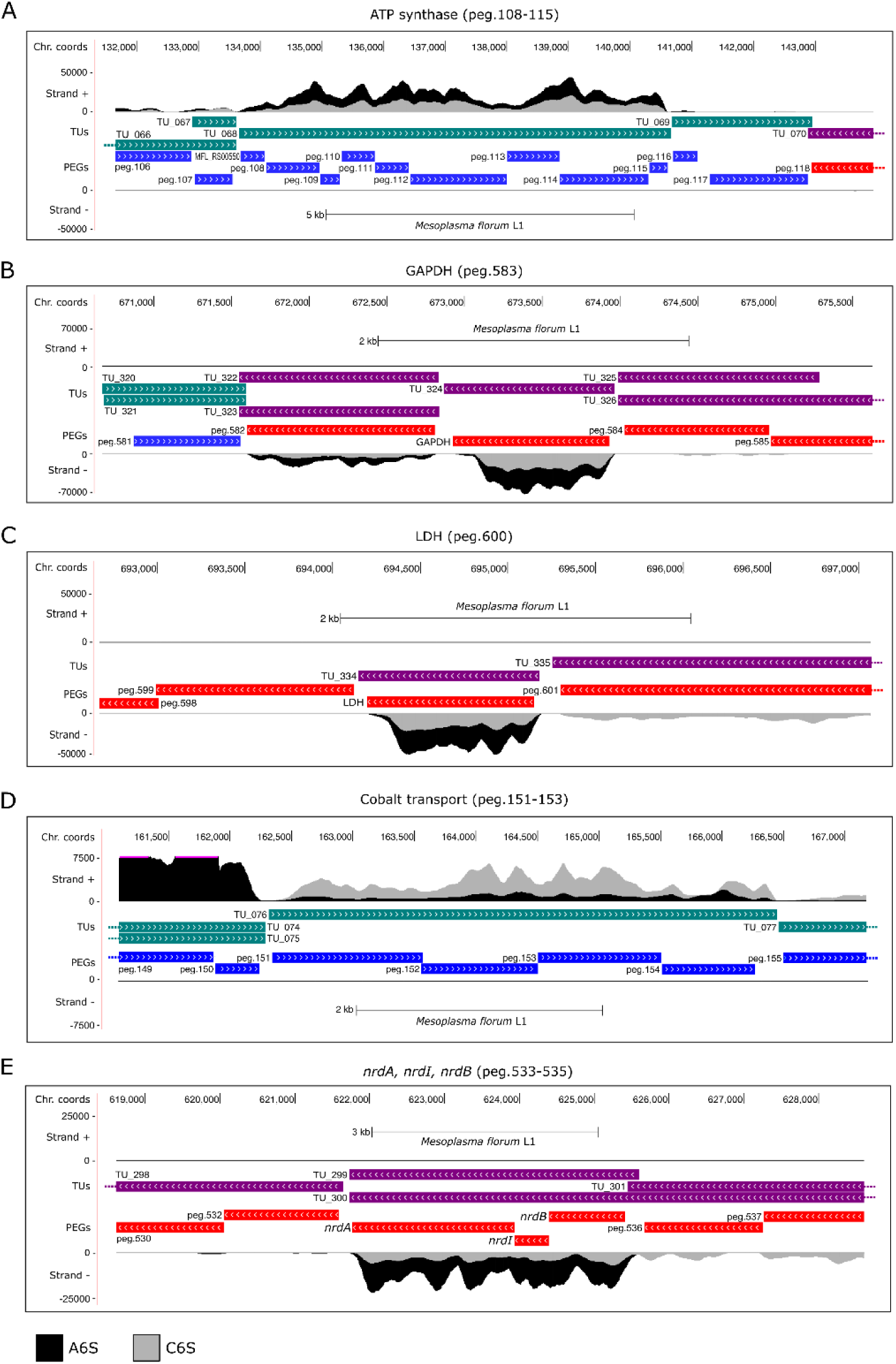
Additional examples of *M. florum* genomic loci showing differential expression in A6S versus C6S. **A)** ATP synthase operon (peg.108-115). **B)** GAPDH encoding gene (peg.583). **C)** LDH encoding gene (peg.600). **D)** *nrdA, nrdI,* and *nrdB* encoding genes (peg.533-535). **E)** Cobalt transport associated genes (peg.151-153). Median strand-specific RNA-seq signal is shown for both conditions (A6S, black; C6S, gray), smoothed over a 16-pixel window. Enhanced RAST annotation is shown, along with previously published *M. florum* transcription units (TU) (Matteau *et al*, 2020). Colored dotted lines indicate genes cropped for representation purposes.

**Figure S10.**
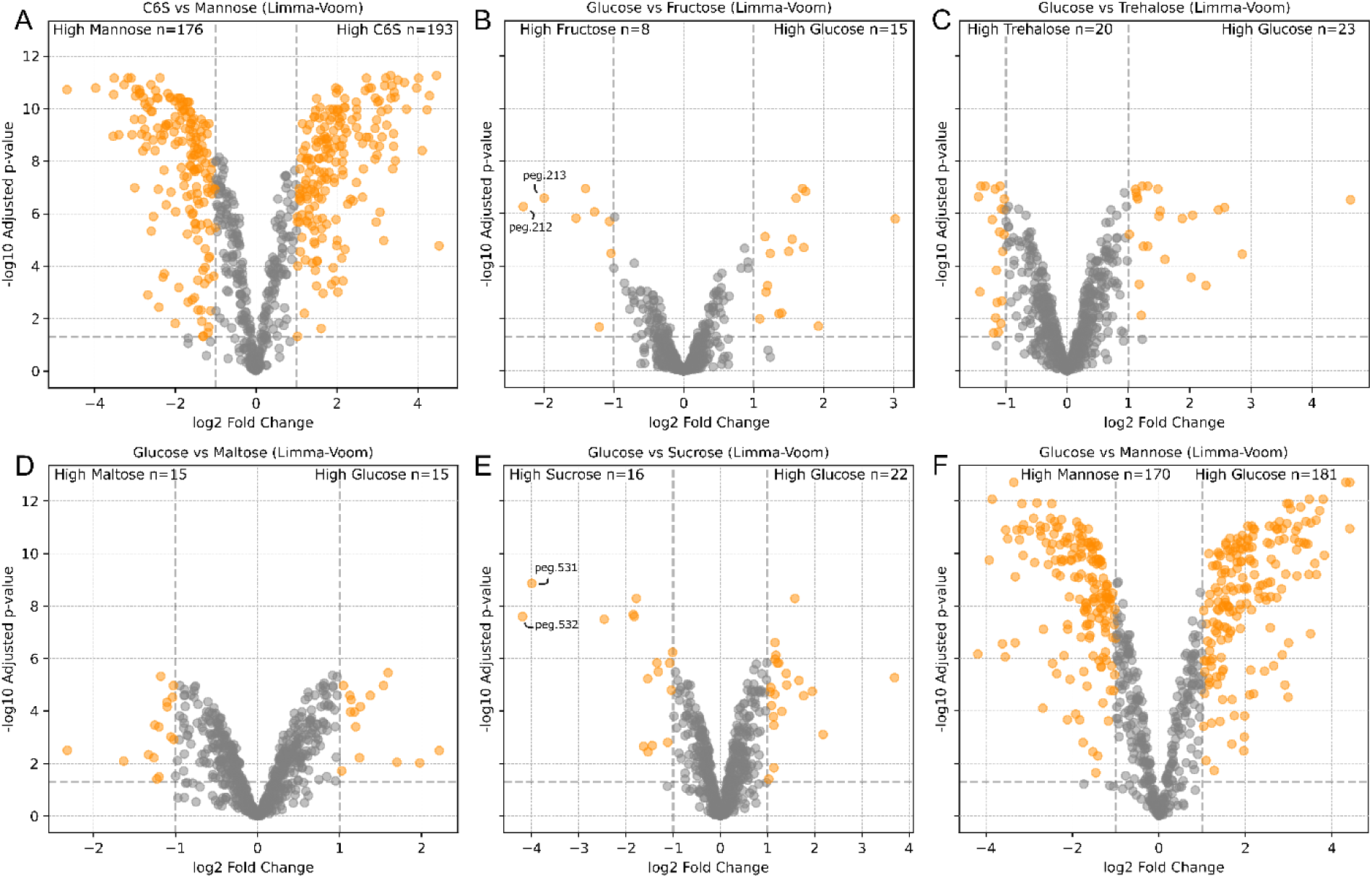
Pairwise differential expression analyses between carbon source conditions. Each point represents a protein-encoding gene (PEG), plotted as log2 fold change (x-axis) versus −log10 adjusted *p*-value (y-axis). Orange points indicate PEGs passing both the fold change (|log2FC| ≥ 1) and adjusted *p*-value (FDR ≤ 0.05) thresholds (gray dashed lines); the number of upregulated genes in each condition is annotated in the corresponding corner. **A)** C6S vs. mannose. **B)** Glucose vs. fructose. **C)** Glucose vs. trehalose. **D)** Glucose vs. maltose. **E)** Glucose vs. sucrose. **F)** Glucose vs. mannose. Differential expression was assessed using the Limma-Voom framework (Law *et al*, 2014).

**Figure S11.**
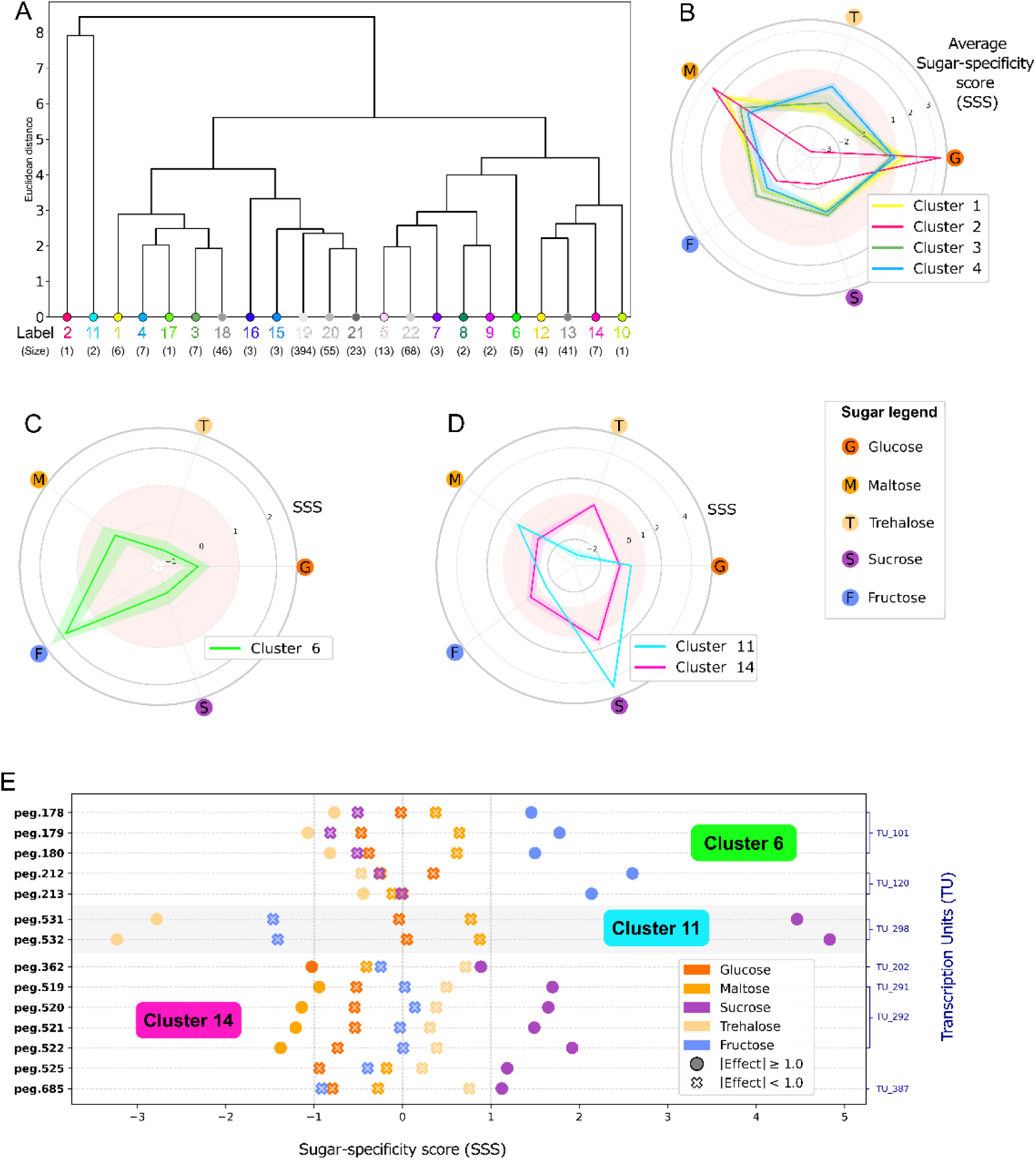
Sugar-specificity score profiles resolve distinct groups of co-regulated genes in *M. florum*. **A)** Complete-linkage hierarchical clustering dendrogram of the 22 clusters, computed on cluster-level SSS profiles. Clusters are colored consistently with panel Fig 5A and cluster size (number of genes) is indicated in parentheses below each label. **B-D)** Radar plots summarizing the mean SSS profile of each selected Group across the five sugar conditions; (B) Group I; Cluster 1-2-3 and 4; (C) Group II; Cluster 6 and (D) Group III; Cluster 11 and 14. The shaded area around the lines represents the standard deviation of each cluster. Axes extend from negative to positive SSS values, with zero at center., with low-significance The pink shaded area from −1 to 1 represents the low-significance section. **E)** Cleveland-style dot plot for Group II and III, illustrating detailed gene-level SSS values for each sugar condition. Each point represents a gene-condition pair; circle markers indicate an effect size ≥ 1.0, and cross markers indicate an effect size < 1.0, distinguishing shifts that are large relative to within-condition dispersion from those that are not. Brackets on the right denote transcription unit (TU) membership (Matteau *et al*, 2020).

**Figure S12.**
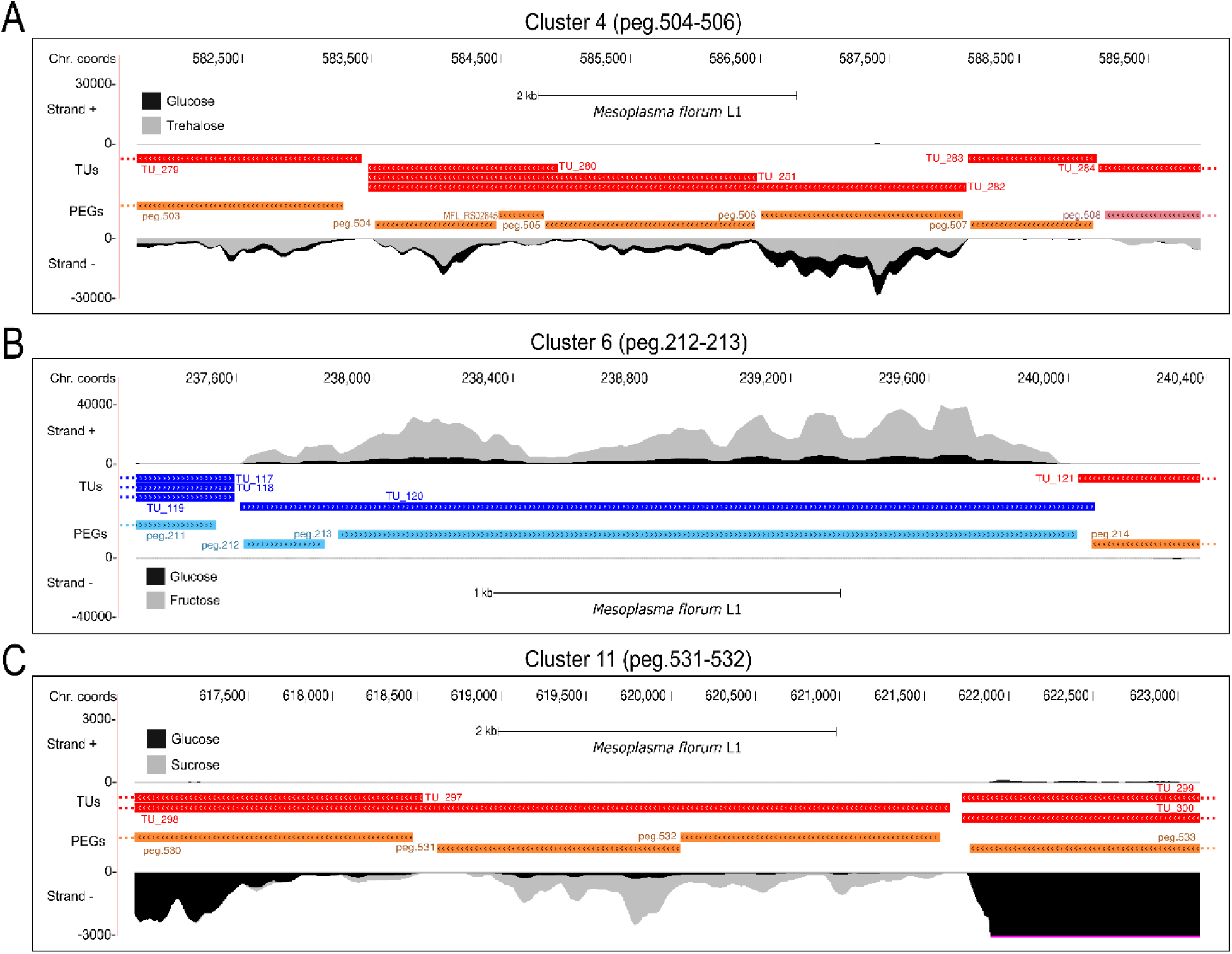
Examples of RNA-seq signal showing differential expression in C6S glucose compared to other sugars. **A)** Cluster 4 (peg.504-506). **B)** Cluster 6 (peg.212-213). **C)** Cluster 11 (peg.531-532). Median strand-specific RNA-seq signal is shown for both conditions (C6S glucose, black; C6S with compared sugars, gray), smoothed over a 16-pixel window. Enhanced RAST annotation is shown, along with previously published *M. florum* transcription units (TU) (Matteau *et al*, 2020). Colored dotted lines indicate genes cropped for representation purposes.

File S1. Original *M. florum* L1 RAST genome annotation file generated in (Baby *et al*, 2018b).

File S2. *M. florum* L1 GenBank 2014 genome annotation file (<u>AE017263.1</u>).

File S3. *M. florum* L1 RefSeq 2022 genome annotation file (<u>NC_006055.1</u>).

File S4. *M. florum* L1 PATRIC 2015 genome annotation file (<u>265311.5</u>).

File S5. *M. florum* L1 enhanced RAST genome annotation file generated in this study.

**Table S1. RNA-seq libraries sequenced in this study.**

**Table S2. Detailed information about GenBank, RefSeq, and PATRIC genome annotations partially overlapping RAST annotations.**

**Table S3. Raw read counts of RNA-seq samples.**

**Table S4. DESeq2 and Lima-Voom normalized read counts and calculated fold changes between selected RNA-seq conditions.**

**Table S5. Functional enrichment analysis of differentially expressed genes between A6S and C6S conditions, grouped into three functional level categories and separated by medium type.** See File S5 for details concerning functional categories.

**Table S6. Functional enrichment analysis of differentially expressed genes between C6S and CMRLAT mannose conditions, grouped into three functional level categories and separated by medium type.** See File S5 for details concerning functional categories.

**Table S7. Oligonucleotides used in this study.**

**Table S8. Expression and clustering metrics for selected genes, and information on selected clusters across five energy source conditions.**

**Table S9. Phosphotransferase system (PTS) analysis, UniProt and InterPro curation.**

## BIBLIOGRAPHY

Baby V, Labroussaa F, Brodeur J, Matteau D, Gourgues G, Lartigue C & Rodrigue S (2018a) Cloning and Transplantation of the *Mesoplasma florum* Genome. ACS Synth Biol 7: 209–217

Baby V, Lachance J-C, Gagnon J, Lucier J-F, Matteau D, Knight T & Rodrigue S (2018b) Inferring the Minimal Genome of *Mesoplasma florum* by Comparative Genomics and Transposon Mutagenesis. mSystems 3: e00198-17

Baby V, Matteau D, Knight TF & Rodrigue S (2013) Complete Genome Sequence of the Mesoplasma florum W37 Strain. Genome Announc 1: e00879–13

Béven L, Charenton C, Dautant A, Bouyssou G, Labroussaa F, Sköllermo A, Persson A, Blanchard A & Sirand-Pugnet P (2012) Specific Evolution of F1-Like ATPases in Mycoplasmas. PLoS ONE 7: e38793

Bosdriesz E, Molenaar D, Teusink B & Bruggeman FJ (2015) How fast-growing bacteria robustly tune their ribosome concentration to approximate growth-rate maximization. FEBS J 282: 2029–2044

Breuer M, Earnest EE, Merryman C, Wise KS, Sun L, Lynott MR, Hutchison CA, Smith HO, Lapek JD, Gonzalez DJ, et al (2019) Essential metabolism for a minimal cell. eLife 8: e36842

Buchweitz LF, Yurkovich JT, Blessing C, Kohler V, Schwarzkopf F, King ZA, Yang L, Jóhannsson F, Sigurjónsson ÓE, Rolfsson Ó, et al (2020) Visualizing metabolic network dynamics through time-series metabolomic data. BMC Bioinformatics 21: 130

Burgos R, Garcia-Ramallo E, Shaw D, Lluch-Senar M & Serrano L (2023) Development of a Serum-Free Medium To Aid Large-Scale Production of *Mycoplasma*-Based Therapies. Microbiol Spectr 11: e04859–22

Casper J, Speir ML, Raney BJ, Perez G, Nassar LR, Lee CM, Hinrichs AS, Gonzalez JN, Fischer C, Diekhans M, et al (2026) The UCSC Genome Browser database: 2026 update. Nucleic Acids Res 54: D1331–D1335

Chen S, Zhou Y, Chen Y & Gu J (2018) fastp: an ultra-fast all-in-one FASTQ preprocessor. Bioinformatics 34: i884–i890

Christodoulou DC, Gorham JM, Herman DS & Seidman JG (2011) Construction of Normalized RNA-seq Libraries for Next-Generation Sequencing Using the Crab Duplex-Specific Nuclease. Curr Protoc Mol Biol 94

Danecek P, Bonfield JK, Liddle J, Marshall J, Ohan V, Pollard MO, Whitwham A, Keane T, McCarthy SA, Davies RM, et al (2021) Twelve years of SAMtools and BCFtools. GigaScience 10: giab008

diCenzo GC, Benedict AB, Fondi M, Walker GC, Finan TM, Mengoni A & Griffitts JS (2018) Robustness encoded across essential and accessory replicons of the ecologically versatile bacterium Sinorhizobium meliloti. PLOS Genet 14: e1007357

Feijó Delgado F, Cermak N, Hecht VC, Son S, Li Y, Knudsen SM, Olcum S, Higgins JM, Chen J, Grover WH, et al (2013) Intracellular Water Exchange for Measuring the Dry Mass, Water Mass and Changes in Chemical Composition of Living Cells. PLoS ONE 8: e67590

Fernandes AD, Reid JN, Macklaim JM, McMurrough TA, Edgell DR & Gloor GB (2014) Unifying the analysis of high-throughput sequencing datasets: characterizing RNA-seq, 16S rRNA gene sequencing and selective growth experiments by compositional data analysis. Microbiome 2: 15

Fu X & Shen Y (2024) Synthetic Genomics: Repurposing Biological Systems for Applications in Engineering Biology. ACS Synth Biol 13: 1394–1399

Gardella RS & Del Giudice RA (1995) Growth of Mycoplasma hyorhinis cultivar alpha on semisynthetic medium. Appl Environ Microbiol 61: 1976–1979

Garrido V, Piñero-Lambea C, Rodriguez-Arce I, Paetzold B, Ferrar T, Weber M, Garcia-Ramallo E, Gallo C, Collantes M, Peñuelas I, et al (2021) Engineering a genome- reduced bacterium to eliminate Staphylococcus aureus biofilms in vivo. Mol Syst Biol 17: MSB202010145

Gaspari E, Malachowski A, Garcia-Morales L, Burgos R, Serrano L, Martins Dos Santos VAP & Suarez-Diez M (2020) Model-driven design allows growth of Mycoplasma pneumoniae on serum-free media. Npj Syst Biol Appl 6: 33

Gibson DG, Glass JI, Lartigue C, Noskov VN, Chuang R-Y, Algire MA, Benders GA, Montague MG, Ma L, Moodie MM, et al (2010) Creation of a Bacterial Cell Controlled by a Chemically Synthesized Genome. Science 329: 52–56

Glass JI, Merryman C, Wise KS, Hutchison CA & Smith HO (2017) Minimal Cells-Real and Imagined. Cold Spring Harb Perspect Biol 9: a023861

Hunter JD (2007) Matplotlib: A 2D Graphics Environment. Comput Sci Eng 9: 90–95

Hutchison CA, Chuang R-Y, Noskov VN, Assad-Garcia N, Deerinck TJ, Ellisman MH, Gill J, Kannan K, Karas BJ, Ma L, et al (2016) Design and synthesis of a minimal bacterial genome. Science 351: aad6253

Jensen CS, Norsigian CJ, Fang X, Nielsen XC, Christensen JJ, Palsson BO & Monk JM (2020) Reconstruction and Validation of a Genome-Scale Metabolic Model of Streptococcus oralis (iCJ415), a Human Commensal and Opportunistic Pathogen. Front Genet 11: 116

Kanehisa M, Sato Y, Kawashima M, Furumichi M & Tanabe M (2016) KEGG as a reference resource for gene and protein annotation. Nucleic Acids Res 44: D457– D462

Karp PD, Paley S, Caspi R, Kothari A, Krummenacker M, Midford PE, Moore LR, Subhraveti P, Gama-Castro S, Tierrafria VH, et al (2025) The EcoCyc database (2025). EcoSal Plus 13: eesp-0019-2024

Karr JR, Sanghvi JC, Macklin DN, Gutschow MV, Jacobs JM, Bolival B, Assad-Garcia N, Glass JI & Covert MW (2012) A Whole-Cell Computational Model Predicts Phenotype from Genotype. Cell 150: 389–401

Lachance J, Matteau D, Brodeur J, Lloyd CJ, Mih N, King ZA, Knight TF, Feist AM, Monk JM, Palsson BO, et al (2021) Genome-scale metabolic modeling reveals key features of a minimal gene set. Mol Syst Biol 17: MSB202010099

Langmead B, Wilks C, Antonescu V & Charles R (2019) Scaling read aligners to hundreds of threads on general-purpose processors. Bioinformatics 35: 421–432

Law CW, Chen Y, Shi W & Smyth GK (2014) voom: precision weights unlock linear model analysis tools for RNA-seq read counts. Genome Biol 15: R29

Lex A, Gehlenborg N, Strobelt H, Vuillemot R & Pfister H (2014) UpSet: Visualization of Intersecting Sets. IEEE Trans Vis Comput Graph 20: 1983–1992

Liao Y, Smyth GK & Shi W (2014) featureCounts: an efficient general purpose program for assigning sequence reads to genomic features. Bioinformatics 30: 923–930

Love MI, Huber W & Anders S (2014) Moderated estimation of fold change and dispersion for RNA-seq data with DESeq2. Genome Biol 15: 550

Matteau D, Duval A, Baby V & Rodrigue S (2024) Mesoplasma florum: a near-minimal model organism for systems and synthetic biology. Front Genet 15: 1346707

Matteau D, Lachance J, Grenier F, Gauthier S, Daubenspeck JM, Dybvig K, Garneau D, Knight TF, Jacques P & Rodrigue S (2020) Integrative characterization of the near- minimal bacterium Mesoplasma florum. Mol Syst Biol 16: MSB20209844

Matteau D, Pepin M-E, Baby V, Gauthier S, Arango Giraldo M, Knight TF & Rodrigue S (2017) Development of *oriC*-Based Plasmids for Mesoplasma florum. Appl Environ Microbiol 83: e03374–16

Matteau D & Rodrigue S (2021) An engineered Mycoplasma pneumoniae to fight Staphylococcus aureus. Mol Syst Biol 17: MSB202110574

McInnes L, Healy J, Saul N & Großberger L (2018) UMAP: Uniform Manifold Approximation and Projection. J Open Source Softw 3: 861

Mölder F, Jablonski KP, Letcher B, Hall MB, Van Dyken PC, Tomkins-Tinch CH, Sochat V, Forster J, Vieira FG, Meesters C, et al (2025) Sustainable data analysis with Snakemake. F1000Research 10: 33

Moore LR, Caspi R, Boyd D, Berkmen M, Mackie A, Paley S & Karp PD (2024) Revisiting the y-ome of *Escherichia coli*. Nucleic Acids Res 52: 12201–12207

Morowitz HJ (1984) The completeness of molecular biology. Isr J Med Sci 20: 750–753

Nordström K, Bagger S, Sillén LG, Kulonen E, Brunvoll J, Bunnenberg E, Djerassi C & Records R (1966) Yeast Growth and Glycerol Formation. Acta Chem Scand 20: 1016–1025

Paysan-Lafosse T, Blum M, Chuguransky S, Grego T, Pinto BL, Salazar GA, Bileschi ML, Bork P, Bridge A, Colwell L, et al (2023) InterPro in 2022. Nucleic Acids Res 51: D418–D427

Pedregosa F, Varoquaux G, Gramfort A, Michel V, Thirion B, Grisel O, Blondel M, Prettenhofer P, Wiess R, Dubourg V, et al (2011) Scikit-learn: Machine Learning in Python. J Mach Learn Res 12: 2825–2830

Python Software Foundation (2020) Python.

Quinlan AR & Hall IM (2010) BEDTools: a flexible suite of utilities for comparing genomic features. Bioinformatics 26: 841–842

R Core Team (2021) R: A language and environment for statistical computing.

Sastry AV, Hu A, Heckmann D, Poudel S, Kavvas E & Palsson BO (2021) Independent component analysis recovers consistent regulatory signals from disparate datasets. PLOS Comput Biol 17: e1008647

Shirvan MH & Rottem S (1993) Ion Pumps and Volume Regulation in Mycoplasma. In Mycoplasma Cell Membranes, Rottem S & Kahane I (eds) pp 261–292. Boston, MA: Springer US

Sirand-Pugnet P, Lartigue C, Marenda M, Jacob D, Barré A, Barbe V, Schenowitz C, Mangenot S, Couloux A, Segurens B, et al (2007) Being Pathogenic, Plastic, and Sexual while Living with a Nearly Minimal Bacterial Genome. PLoS Genet 3: e75

Srinivasan M, Scheinost JC, Petela NJ, Gligoris TG, Wissler M, Ogushi S, Collier JE, Voulgaris M, Kurze A, Chan K-L, et al (2018) The Cohesin Ring Uses Its Hinge to Organize DNA Using Non-topological as well as Topological Mechanisms. Cell 173: 1508–1519.e18

Talenton V, Baby V, Gourgues G, Mouden C, Claverol S, Vashee S, Blanchard A, Labroussaa F, Jores J, Arfi Y, et al (2022) Genome Engineering of the Fast-Growing *Mycoplasma feriruminatoris* toward a Live Vaccine Chassis. ACS Synth Biol 11: 1919–1930

The UniProt Consortium, Bateman A, Martin M-J, Orchard S, Magrane M, Ahmad S, Alpi E, Bowler-Barnett EH, Britto R, Bye-A-Jee H, et al (2023) UniProt: the Universal Protein Knowledgebase in 2023. Nucleic Acids Res 51: D523–D531

trackhub (2017)

Virtanen P, Gommers R, Oliphant TE, Haberland M, Reddy T, Cournapeau D, Burovski E, Peterson P, Weckesser W, Bright J, et al (2020) SciPy 1.0: fundamental algorithms for scientific computing in Python. Nat Methods 17: 261–272

Waskom M (2021) seaborn: statistical data visualization. J Open Source Softw 6: 3021

Zharova TV, Grivennikova VG & Borisov VB (2023) F1·Fo ATP Synthase/ATPase: Contemporary View on Unidirectional Catalysis. Int J Mol Sci 24: 5417

